# Organic electron donors and terminal electron acceptors structure anaerobic microbial communities and interactions in a permanently stratified sulfidic lake

**DOI:** 10.1101/2021.02.16.431432

**Authors:** Connie A. Rojas, Ana De Santiago Torio, Serry Park, Tanja Bosak, Vanja Klepac-Ceraj

## Abstract

The extent to which nutrients structure microbial communities in permanently stratified lakes is not well understood. This study characterized microbial communities from the anoxic layers of the meromictic and sulfidic Fayetteville Green Lake (FGL), NY, and investigated the roles of organic electron donors and terminal electron acceptors in shaping microbial community structure and interactions. Bacterial communities from the permanently stratified layer below the chemocline (monimolimnion) and from enrichment cultures inoculated by lake sediments were analyzed using 16S rRNA gene sequencing. Results showed that anoxygenic phototrophs dominated microbial communities in the upper monimolimnion (21 m), which harbored little diversity, whereas the most diverse communities resided at the bottom of the lake (~52 m). Organic electron donors explained 54% of the variation in the microbial community structure in aphotic cultures enriched on an array of organic electron donors and different inorganic electron acceptors. Electron acceptors only explained 10% of the variation, but were stronger drivers of community assembly in enrichment cultures supplemented with acetate or butyrate compared to the cultures amended by chitin, lignin or cellulose. We identified a range of habitat generalists and habitat specialists in both the water column and enrichment samples using Levin’s index. Network analyses of interactions among microbial groups revealed Chlorobi and sulfate reducers as central to microbial interactions in the upper monimolimnion, while Syntrophaceae and other fermenting organisms were more important in the lower monimolimnion. The presence of photosynthetic microbes and communities that degrade chitin and cellulose much below the chemocline supported the downward transport of microbes, organic matter and oxidants from the surface and the chemocline. Collectively, our data suggest niche partitioning of bacterial communities by interactions that depend on the availability of different organic electron donors and terminal electron acceptors. Thus, light, as well as the diversity and availability of chemical resources drive community structure and function in FGL, and likely in other stratified, meromictic lakes.

## 1 INTRODUCTION

Microbial communities in permanently stratified ecosystems, such as sulfidic meromictic lakes, mediate the biogeochemical cycling of carbon, oxygen, sulfur and other elements (Bosshard et al., 2000; Lehours et al., 2005; Lehours et al., 2007; Lauro et al., 2011; Comeau et al., 2012; Baricz et al., 2014; Andrei et al., 2015; Baatar et al., 2016; Meyerhof et al., 2016). A steep chemical gradient known as the chemocline separates layers of oxic and anoxic waters and creates distinct niches for bacterial metabolisms and interspecies interactions. For example, oxygenic photosynthesis and aerobic respiration prevail in the oxygenated surface layer, while anoxygenic phototrophs are common at or immediately below the chemocline. In even deeper waters, photosynthetic bacteria are replaced by sulfur-cycling chemotrophs and fermenting microorganisms. The structure of microbial communities in several meromictic lakes has been described (Humayoun et al., 2003; Lehours et al., 2005; Dimitriu et al., 2008; Klepac-Ceraj et al., 2012; Baatar et al., 2016; Hamilton et al., 2016; Danza et al., 2017), but much less is known about the biotic and abiotic factors that govern the assembly of these communities and the biologically-mediated coupling of elemental cycles.

Here, we sought to identify relationships between the structure of microbial communities and biogeochemical cycling in Fayetteville Green Lake (FGL, **Figure 1**). The lake is located in upstate New York, NY, USA (Eggleton, 1931; Deevey et al., 1963; Brunskill and Ludlam, 1969; Hilfinger IV et al., 2001; Zerkle et al., 2010; Havig et al., 2015) and is an analog for understanding the sulfide-rich, oxygen-depleted Proterozoic oceans (Canfield, 1998; Lyons et al., 2009), modern sulfidic environments (Arthur et al., 1988; Kuypers et al., 2001; Johnston et al., 2005; Nabbefeld et al., 2010), and the cycling of metals such as iron and manganese in the presence of sulfide (Torgersen et al., 1981; Havig et al., 2015). Small and moderately deep, the lake reaches 53 meters at its deepest point and has a thick chemocline between 15 and 21 meters (Eggleton, 1956; Brunskill and Ludlam, 1969; Torgersen et al., 1981; Havig et al., 2015). The chemocline is maintained by the input of sulfate- and calcium-rich groundwater that flows through the gypsum-rich Vernon Shale Formation (Brunskill and Ludlam, 1969). The anoxic deep waters contain high concentrations of sulfide (up to 1.86 mM) and sulfate (between ~11.7 and 15.1 mM) (Brunskill and Ludlam, 1969; Torgersen et al., 1981; Hilfinger IV et al., 2001; Zerkle et al., 2010; Meyer et al., 2011; Havig et al., 2015). The concentrations of sulfide, methane (CH_4_) and dissolved inorganic carbon (DIC) increase with depth below the chemocline and vary little seasonally. The increase of methane concentrations coincides with a bacterial plate located at the base of the chemocline around 20 m and reaches a maximum value of 23.5 μM in the lower monimolimnion (Havig et al., 2018). The negative δ^13^C values of methane (~ −100‰) suggest biogenic production. Additional geochemical indicators (Table S1, Figure 1; Fry, 1986; Zerkle et al., 2010) suggest the presence of a second chemocline at 45 m. Fe(II) and Mn(II) concentrations peak within the chemocline and their maxima coincide with a dense microbial plate (Torgersen et al., 1981; Havig et al., 2015). The concentration of dissolved organic carbon (DOC) of ~ 200 μM and DOC δ^13^C values of ~ −31‰ throughout the entire water column suggest rapid recycling of labile DOC and the accumulation of recalcitrant organic carbon with depth (Havig et al., 2015; Havig et al., 2018). All these observations urge a better understanding of microbial community structure and function across different depths in FGL. Furthermore, because the activity of microbial communities may influence the accumulation and sequestration of DOC, research on the nutrient cycling and the predicted responses of communities in euxinic lakes to the changing climate deserve more attention (Anderson et al., 2013; Anderson et al., 2014).

**Figure 1.**
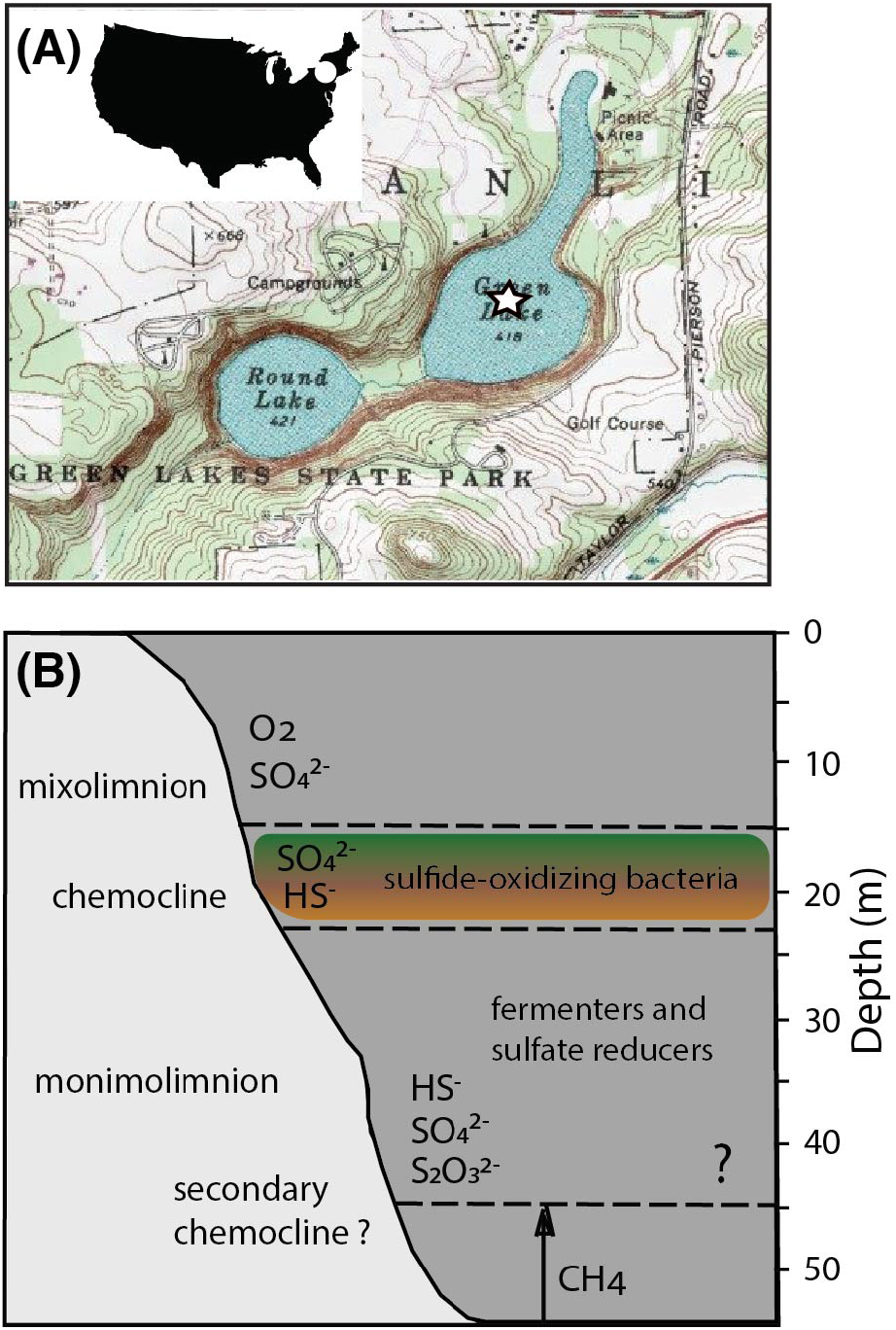
Location and biogeochemical profiles of FGL. (A) U.S. Geological Survey map of FGL and the surrounding area. The star indicates the sampling site at the deepest part of the lake. (B) Stratification of major microbial metabolisms, O_2_, H_2_S and methane in FGL. Question mark indicates the depth of the hypothesized secondary chemocline (Zerkle et al., 2010).

The biological diversity of FGL has been characterized primarily in the photic zone. Green algae, zooplankton and unicellular phototrophic cyanobacteria inhabit the upper, oxygen-rich layer (Culver and Brunskill, 1969). In this layer, cyanobacteria (*Synechococcus* sp.) seasonally stimulate the precipitation of calcium carbonate and cause “whiting events” by forming clumps of CaCO_3_ (Thompson and Ferris, 1990; Thompson et al., 1997). These clumps deliver organic matter and calcite to the sediments and contribute to the light-colored, calcite-rich and dark-colored, organic-rich laminae (Brunskill, 1969; Ludlam, 1969; Hubeny et al., 2011). At the chemocline, purple and green anoxygenic photosynthetic sulfide-oxidizing bacteria form a dense layer and are responsible for 83% of the annual primary production in the lake (Culver and Brunskill, 1969; Meyer et al., 2011). Photosynthetic pigments produced by anoxygenic phototrophs have also been detected in the sediments (Meyer et al., 2011; Fulton et al., 2018), attesting to the delivery of photosynthetic microbes from the chemocline to the deeper parts of the water column.

Few studies have investigated microbial activity and interactions below the chemocline and their contributions to biogeochemical cycling in meromictic lakes such as FGL. Metagenomic reconstructions of the most abundant genes in other meromictic lakes suggest that sulfate reduction and fermentation are central to the cycling of carbon and sulfur in the lower anoxic zone (Biderre-Petit et al., 2011; Gies et al., 2014; Llorens-Mares et al., 2015; Hamilton et al., 2016). However, the factors that shape microbial diversity and microbial interactions in the aphotic zone of FGL and other euxinic lakes remain poorly understood. Here, we use a combination of 16S rRNA gene sequencing, enrichment cultures and network analyses to: 1) characterize the composition and diversity of bacterial communities below the chemocline in FGL; 2) determine how organic electron donors and terminal electron acceptors drive microbial community structure and function below the photic zone; and 3) elucidate key microbial interactions potentially involved in nutrient cycling in meromictic lakes. Comparisons of the 16S rRNA gene surveys of microbial communities sampled from different depths of the lake along with those obtained under a range of culturing conditions reveal specific microbe-microbe interactions that may couple and mediate the biogeochemical cycling of carbon and other elements in meromictic lakes.

## 2 MATERIALS AND METHODS

### 2.1 Water column sampling

Samples from Fayetteville Green Lake, Fayetteville, New York, U.S.A (43°3′N, 75°57.6″ W) were collected on August 14, 2013. We used a 12 L Niskin bottle to collect water samples in triplicates from five different depths (21 m, 24 m, 30 m, 45 m, and 52 m) and a grab corer to collect samples from the underlying sediments (~53 m). Methane concentrations, pH values and temperatures measured at the time of sampling are provided in **Table S1**. A permit to conduct sampling at FGL was obtained from the New York State Office of Parks, Recreation and Historic Preservation. Upon collection, all water samples were kept on ice for approximately 6 hours and filtered through a 0.22 μm Sterivex GV filter (Millipore) using a Masterflex L/S 7553-70 peristaltic pump. Both filter samples and sediments were frozen at −80 °C until DNA extraction. The anaerobic sediment slurry was kept at 4 °C in the dark in tightly capped glass jars or serum bottles for later inoculations. **Table S2** presents all water column samples used in this study and their corresponding depths.

### 2.2 Enrichment cultures

To better understand how terminal electron acceptors and organic electron donors shape microbial community composition, we inoculated 3 mL of lake sediment slurry into 97 mL of Green Lake medium (GL) supplemented with different terminal electron acceptors and organic electron donors. The recipe for the Green Lake (GL) medium omitted the dominant ions to experimentally control for electron acceptors, but otherwise, the recipe reflects the background ion concentrations in the lake as reported by Brunskill and Ludlam (1969). All chemicals were purchased from Sigma-Aldrich (St. Louis, MO). The basal GL medium contained: KH_2_PO_4_ 13.6 mg/L, NH_4_Cl 0.3 g/L, KCl 67.1 mg/L, and NaHCO_3_ 9 g/L. Trace metal solution SL-10 (1000x) was composed of HCl (25%; 7.7 M) 10 mL, FeCl_2_ × 4 H_2_O 1.5 g/L, ZnCl_2_ 70 mg/L, MnCl_2_ x 4 H_2_O 100 mg/L, H_3_BO_3_ 6 mg/L, CoCl_2_ x 6 H_2_O 190 mg/L, CuCl_2_ x 2 H_2_O 2 mg/L, Na_2_MoO_4_ × 2 H_2_O 31mg/L. The vitamin solution (100x) contained biotin 2 mg/L, folic acid 2 mg/L, pyridoxine-HCl 10 mg/L, thiamine-HCl x 2 H_2_O 5 mg/L, riboflavin 5 mg/L, nicotinic acid 5 mg/L, D-Ca-pantothenate 5 mg/L, vitamin B12 0.1 mg/L, p-aminobenzoic acid 5 mg/L, lipoic acid 5 mg/L. Filter-sterilized vitamin solution (10 mL) and trace element solution SL-10 (1 mL) were added to the autoclaved GL medium to a final volume of 1 L. After autoclaving, the medium was cooled under an 80% N_2_, 20% CO_2_ gas mixture and flushed with N_2_. The final pH of the medium was 7.2.

Terminal electron acceptors (FeCl_3_, Na_2_SO_4_, S^0^) and organic electron donors (acetate, propionate, butyrate, lignin, chitin, cellulose) were sterilized separately and added to the medium. Each enrichment culture was supplemented with one of the three terminal electron acceptors and one of the seven electron donors or no carbon (NoC). Acetate, butyrate, propionate and sulfate were prepared as anaerobic 1 M stock solutions and their final concentrations in enrichments were 10 mM. Fe(III) was added as poorly crystalline ferrihydrite prepared to the final concentration of 10 mM ferrihydrite and titrated with 10 N NaOH to pH 7 as described in Shang et al. (2020). Elemental sulfur (MW=32.08 g/mol) was heat sterilized at 110°C overnight and added to a bottle containing FGL medium to generate a ~1 M stock solution. This solution was mixed by shaking and used in the enrichments at the final concentration of ~10 mM. Individual stock solutions of chitin (from shrimp shells, MW=203.19 g/mol), lignin (MW=770.9 g/mol), and cellulose (MW=162.14 g/mol) were prepared by adding the dry chemicals to FGL medium to 1M, autoclaving and flushing the headspace with an 80% N_2_, 20% CO_2_ gas mixture. Cellulose, lignin, and chitin are not very soluble, so their stock solutions were mixed by shaking before adding aliquots to the GL medium to a final concentration of ~10 mM. Methane was introduced into enrichment cultures by flushing the headspace (25 mL) with a 5% CH_4_, 15% CO_2_ and 80% N_2_ gas mixture. The final concentration of CH_4_ in solution likely did not exceed 1 mM, and was well above the reported CH_4_ concentration of FGL (Havig et al., 2018).

To establish the original enrichment, 5 mL of sediment slurry (1 part sediment to 1 part GL medium) was transferred into 95 mL of FGL medium amended with one electron acceptor and one electron donor. Two additional transfers used 5 mL of previous generation as the inoculum. Havig et al. (2018) measured ~2% C dry wt of organic carbon in the top 10 cm of the sediment core, and ~8 mM of DOC and ~1 mM of DIC in the top 8 cm of the sediment core. Based on these measurements, we estimated that less than 0.3 mg C dry wt, 62.5 nM of DIC, and 0.5 μM of DOC were present in the final enrichment. All original cultures and subsequent enrichment cultures (100 mL) were grown in duplicates at room temperature in the dark and transferred every four weeks. They were prepared in 125 mL serum bottles capped with butyl rubber stoppers and reduced with 1 mM sulfide. Following the growth of the second transfer from the establishment of the original enrichment from lake sediments, the cultures were harvested to profile the composition of microbial communities by filtering 100 mL of each culture through a 0.22 μm Sterivex GV filter (Millipore) in the anaerobic hood under an 80% N_2_ and 20% CO_2_ gas mixture. The filters with cells were frozen at −80°C until DNA extraction.

All enrichment cultures were grown in the dark to mirror the conditions below the photic zone. The organic additives and methane were selected to reflect the diversity of possible carbon sources in FGL: cellulose and lignin from the plant-derived material, chitin from copepods (Culver and Brunskill, 1969), butyrate, propionate and acetate as products of fermentation and methane as the product of methanogenesis. The terminal electron acceptors represented the most common electron acceptors available in this zone (**Table S1**). See **Table S3** for a list of samples from enrichment cultures and their conditions.

### 2.3 Extraction of DNA from water column samples and enrichment cultures

DNA from Sterivex filters was extracted using the PowerWater Sterivex DNA Isolation Kit^TM^ (MO BIO Laboratories, Carlsbad, CA) following a modified manufacturer recommended protocol. Briefly, instead of adding the lysis reagents directly to the filters, we transferred the filter and its contents into a 5-mL polypropylene snap-cap tube (Eppendorf, Hamburg, Germany) because some filter units contained sample water and could not fit the lysis reagents. Following the transfer, the samples were processed according to the manufacturer’s protocol and eluted in 50 μl of elution buffer. Gel electrophoresis confirmed that sample extractions yielded amplifiable 16S rRNA and negative control (blank) extractions did not. DNA concentrations in all samples and negative controls were quantified using a NanoDrop ND-1000 (Thermo Scientific, Inc., Wilmington, DE, USA). The DNA extracts were stored at −80°C until sequenced.

### 2.4 Amplification and sequencing of 16S rRNA genes

The V3-V4 region of the 16S rRNA gene from samples of the water column was sequenced according to the protocol described by Caporaso and colleagues (2011). This region was amplified using the universal bacterial primers 341F 5’-CCTACGGGAGGCAGCAG-3’ (Muyzer et al., 1993) and 806R 5’-GGACTACHVGGGTWTCTAAT-3 (Caporaso et al., 2011). The appropriate Illumina adapters were included on both primers and the error-correcting 12-bp barcode unique to each sample was included on the reverse primer. Samples were purified using AMPure beads, pooled into a library (100 ng) and quantified by qPCR. Twenty percent of denatured PhiX was added to the amplicon pool (12 pM) and sequenced on the MiSeq platform (Illumina, San Diego, CA) at the Forsyth Institute (Cambridge, MA).

Samples of enrichment cultures were sequenced at the Environmental Sample Preparation and Sequencing Facility at Argonne National Laboratory (Lemont, IL) using the same chemistry and sequencing platform and following the same protocol as described above (Caporaso et al., 2011). The primers targeted the V4 region and used the same reverse primer (806R), but a different forward primer 515F (5’- GTGCCAGCMGCCGCGGTAA-3’). Because of the differences in how the enrichment and lake samples were sequenced, we did not combine the two datasets, but rather characterized each dataset individually and drew inferences between them.

### 2.5 16S rRNA gene sequence processing

For water column samples, paired-end reads were joined using Flash software (Magoc and Salzberg, 2011). Libraries were demultiplexed and reads were filtered using Quantitative Insights into Microbial Ecology (QIIME), v1.9.0 (Caporaso et al., 2010; Caporaso et al., 2012). Any reads that did not assemble by being perfectly matched in the overlapping region or that failed to meet the q-score threshold of 25 were removed from subsequent analyses. At this step, the sequences were imported into QIIME2, v.2020.8 (Bolyen et al., 2019). Chimeric sequences were identified and removed using UCHIME’s usearch61 *de novo* based chimera detection algorithm (Edgar et al., 2011). The quality-filtered sequences were then clustered into operational taxonomic units (OTUs) at 97% sequence identity. Taxonomy was assigned using a Naïve-Bayes classifier compared against the SILVA v.138 reference database trained on the 515F-806R region of the 16S rRNA gene (Bokulich et al., 2018). After quality filtering and the removal of chimeras, each sample had more than 37,500 sequences that were ~ 450 bp long.

Raw sequence reads of samples from enrichment cultures were denoised, filtered and clustered into amplicon sequence variants (ASVs) using the Divisive Amplicon Denoising Algorithm (DADA2) plugin in QIIME 2 v.2020.8 (Callahan et al., 2016). Taxonomy was assigned to each ASV using a Naïve-Bayes classifier compared against SILVA v.138 reference database trained on the 515F-806R region of the 16S rRNA gene (Bokulich et al., 2018).

### 2.6 Statistical analyses of amplicon sequences

We performed most statistical analyses and visualizations using the QIIME2 software package and R statistical program v3.6.2 (R_Core_Team, 2019). To visualize the community composition, we constructed bubble plots and stacked bar plots of bacterial relative abundances using the ggplot2 package (Wickham, 2016). Alpha-diversity was estimated using the Chao 1 richness index and Shannon diversity index (Shannon, 1948; Chao, 1984). To evaluate the influence of depth, organic electron donor, or terminal electron acceptor on the richness and evenness of microbial communities, we constructed linear models in R v3.6.2 (R_Core_Team, 2019). To test whether variables of interest significantly predicted alpha-diversity, we conducted likelihood ratio tests (LRT; α=0.05) on our linear models using the R car package (R_Core_Team, 2019). These were followed by multiple comparison testing using Tukey HSD contrasts and Benjamini-Hochberg adjusted p-values (Benjamini and Hochberg, 1995). Boxplots of microbiota alpha-diversity were generated in ggplot2.

Community structure (i.e., β-diversity) was assessed using Bray-Curtis dissimilarity indices in R. To determine whether variables of interest including sample depth, organic electron donor or terminal electron acceptor shaped microbial community structure, we used permutational multivariate analysis of variance tests (PERMANOVA) (Anderson, 2000) using the adonis function from the R vegan package (Oksanen et al., 2016). For water column data, the model included depth as a single predictor variable, and for enrichment data, the model specified carbon source, electron acceptor, and their interaction as predictors. The clustering of bacterial communities was visualized via principal coordinate analysis (PCoA) plots constructed from Bray-Curtis distance matrices using ggplot2 (Wickham, 2016). To evaluate whether FGL environmental gradients (**Table S1**) were correlated with bacterial community structure, these data were also analyzed using constrained correspondence analysis (CCA) using the cca function from the R vegan package (Oksanen et al., 2016).

The niche breadth for bacterial taxa from both enrichment cultures and water column samples was calculated using Levins’ measure of niche breadth (Levins, 1968). Levins’ breadth is useful for identifying bacterial taxa that may be habitat generalists or specialists. To calculate Levins’ niche breadth index, we measured the uniformity of distribution of individual taxa among lake or enrichment samples using the following:

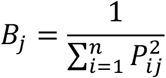

where *Bj* indicates the breadth index of a taxon *j* and *Pij* is the relative abundance of taxon *j* in a given habitat *i*. The habitats consisted of 5 lake depths (**Table S2**) and 24 enrichments (**Table S3**). Taxa with average relative abundances across samples < 2 × 10^−5^ were removed from the analysis to minimize the number of false specialists (Pandit et al., 2009). Bacterial taxa with niche breadths outside of one standard deviation from the mean niche breadth were classified as habitat and resource specialists or generalists (Logares et al., 2013). Scatterplots of Levin’s niche breadth versus average relative abundance for each taxon were plotted in R using the ggplot2 package.

### 2.7 Network construction

To visualize connections between bacterial groups in the water column and in enrichments, we constructed co-occurrence networks using CoNet app available in Cytoscape v3.7.2 (Faust and Raes, 2016). Network inferences were built using taxon abundance tables from 16S rRNA gene profiles. Taxa with fewer than 20 counts across all samples and present in < 3 samples were removed from the dataset. The settings were the same for lake and enrichment datasets. For water column data, networks were generated separately for data from the upper monimolimnion (i.e., shallower water column) and the lower monimolimnion (i.e., deeper waters). For enrichment data, networks were constructed separately for cultures amended with each of the three terminal electron acceptors, S^0^, SO_4_^2−^, Fe(III). To minimize sparsity issues and false correlations due to the zero-inflated data typical of microbiome studies, we filtered for OTU occurrence across samples according to the minimum value suggested by the program (Faust and Raes, 2016).

Co-occurrence and co-exclusion interactions were calculated in CoNet using five measures: Pearson (Pearson, 1895), Spearman (Sedgwick, 2014), Bray-Curtis dissimilarity (Bray and Curtis, 1957), mutual information, and Kullback-Leibler divergence (Kullback and Leibler, 1951). Briefly, the initial top and bottom edge numbers were set at 1000. One thousand permutation scores and 1000 bootstrap scores were computed for each edge and each measure of association. The vectors of taxon pairs were shuffled to minimize any biases associated with composition. The correlation measure-specific p-values were merged using Brown’s method (Brown, 1975) and corrected using the Benjamini-Hochberg method (Benjamini and Hochberg, 1995). Significant interactions (p-value < 0.05) were visualized in Cytoscape (Shannon et al., 2003) and taxa without any significant interactions were removed from the network. Taxa were shown as “nodes” in the networks, where the edges represented significant co-occurrence or co-exclusion relationships. The color of the edges reflected directionality (green: co-occurrence, red: co-exclusion), and their transparency was proportional to the calculated significance of the relationship.

### 2.8 Metagenome sequencing and analysis

Because the primer sets that target the V3–V5 region of the 16S rRNA gene underestimate the abundances of archaeal sequences in biological samples (Pinto and Raskin, 2012), we sequenced metagenomes from the lower monimolimnion, at 45.5 m and 52 m. The metagenome data were used solely to “spot check” amplicon bias in the 16S rRNA gene profiles. The DNA extracts were sent for metagenomic sequencing to the University of Southern California’s Genome and Cytometry Core Facility (Los Angeles, CA, USA). Before sequencing on the Illumina HiSeq 2500 platform, DNA was sheared using dsDNA Shearas Plus (Zymo, Irvine, CA, USA), and cleaned using Agencourt AMPure XP beads (Beckman-Coulter, Indianapolis, IN, USA). The library was quantified using the Qubit 2.0 Fluorometer and the DNA fragment size was determined with an Agilent Bioanalyzer 2100. Raw data were uploaded to MG-RAST, a metagenomics service for analysis of microbial community structure and function (Keegan et al., 2016), to obtain taxonomic profiles for each sample. Raw metagenome sequence files and contigs are available upon request.

## 3 RESULTS

### 3.1 Diversity and structure of bacterial taxa in the FGL monimolimnion

To examine the relationships between *in situ* bacterial communities and the biogeochemical profiles below the chemocline of FGL, we sequenced the 16S rRNA genes of samples from the upper monimolimnion (21 m, 24 m) and the lower monimolimnion (30 m, 45 m, and 52.5 m). Rarefaction analysis indicated that the depths from the upper monimolimnion were more completely sampled than the deeper depths (**Figure S1**). Overall, bacterial communities were almost identical among the replicates, suggesting a high reproducibility of our sampling, sample processing and sequencing methods (**Figure S1**). The only exception was one less diverse sample from 45 m that grouped outside of the other two samples from 45 m (**Figure S1B**). Of the 16S rRNA gene sequences examined, 96.6% were assigned to Bacteria and only 3.4% to Archaea. Although the primer bias against Archaea is known (Pinto and Raskin, 2012), these values closely resembled the values observed in the FGL metagenome data generated to check for the primer bias in the 16S rRNA amplicon data. In the metagenomes, 6.2% (45.5 m) and 5.5% (52.5%) of sequences belonged to Archaea, compared to the 4.5% of sequences recovered from 16S rRNA gene data at the same depths.

Across all sampled depths, the four most dominant bacterial phyla included Bacteroidota (35% average relative abundance), Desulfobacterota (29%; previously characterized as Deltaproteobacteria within the phylum Proteobacteria), Verrucomicrobiota (5%), and Patescibacteria (5%) (**Figure S2A, Table S4**). Collectively, these phyla constituted over 75% of the community in the water column and were detected across all depths. Additional phyla with average relative abundances >1% included Chloroflexi, Campilobacterota (previously part of Epsilonproteobacteria within the phylum Proteobacteria), Nanoarchaeota, Cyanobacteria, Firmicutes, Proteobacteria and Actinobacteriota (previously Actinobacteria) (**Figure S2A, Table S4**). Chlorobia (Chlorobi), Syntrophia (previously classified within Deltraproteobacteria as Synthrophobacteriales), Bacteroidia (Bacteroidetes), and Parcubacteria (OD1) were the four most dominant classes and constituted > 63% of sequences from the water column(**Figure 2A**).

**Figure 2.**
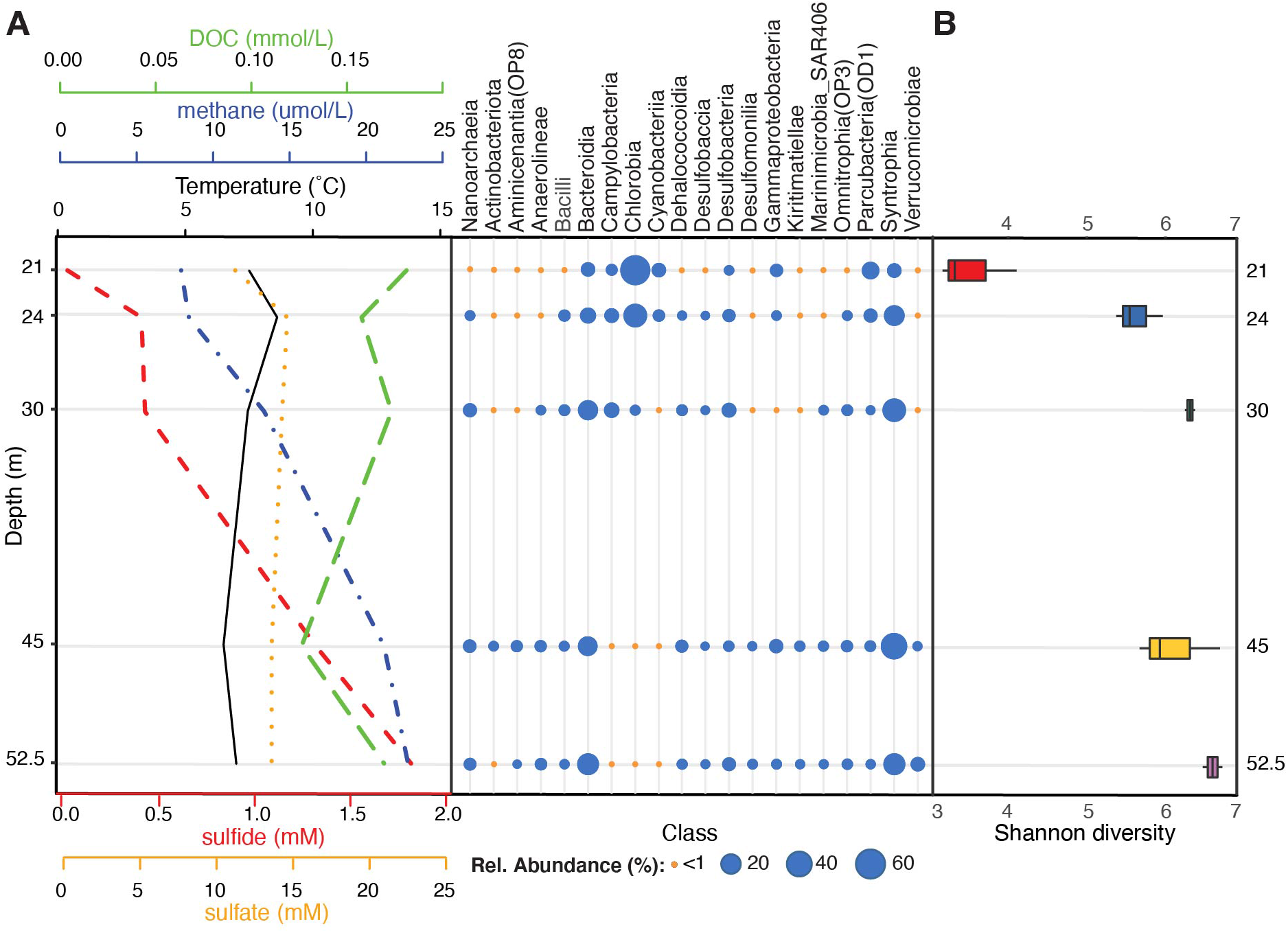
Stratification of bacterial communities below the FGL chemocline. (A) Taxonomic profiles of the mean relative abundances of bacterial classes (SILVA v.138 taxonomy) for each sampled depth (y-axis), alongside sulfide, sulfate, DOC, methane, and temperature profiles. Bacterial classes with at least one depth with >1% mean relative abundance across samples are plotted. Temperature and methane concentrations were measured during sampling in July 2013 while sulfide, sulfate, and DOC profiles were collected by Havig et al. (2015) at the same time (see **Table S1** for values). (B) Microbial community α-diversity (Shannon diversity) at each sampled depth.

A comparison of the overall alpha-diversity between sampled depths revealed an increase in microbial community evenness with depth (**Figure 2C**). Mean Shannon diversity values were 3.18 ± 0.80 for 21 m to 5.96 ± 0.14 for 52.5 m (linear model LRT: Chao 1 Richness χ_4_^2^ =2.31, p=0.14; Shannon diversity χ_4_^2^ =7.24, p=0.009; see **Table S5** for post-hoc comparisons). Within the upper monimolimnion, bacterial communities were dominated by Chlorobia (Chlorobiales) (average relative abundance of 57.30% at 21 m and 29.49% at 24 m). These organisms were much less abundant at 30 m, 45 m and 52 m (average relative abundances of 2.19%, 0.09% and 0.34%, respectively; **Figure 2B**, **Table S4**). ASVs assigned to sulfate reducing Syntrophales (Syntrophia), Sphingobacteriales (Bacteroidia), Woesearchaeales (Nanoarchaeia), and Desulfatiglandales (Desulfobacteria) increased with depth and these ASVs constituted 32.12%, 15.85%, 4.75% and 4.03% of the community below the chemocline, respectively (**Figure 2B** **and S2B)**. Communities at 30 m contained greater abundances of sulfur-oxidizing Campylobacterota (primarily *Sulfurimonas* and *Sulfuricurvum* spp., 8.4%) relative to all other depths, whereas the order Chthoniobacterales had the highest relative abundance at 52 m (Verrucomicrobiae; 6.3%). These organisms constituted less than <1% of the amplicon sequences at all other depths (**Table S4**).

### 3.2 Depth-stratified bacterial community structure and biogeochemistry

Bacterial communities below the chemocline were strongly partitioned by depth: this variable accounted for 35% of the observed variation in community structure across samples (**Figure S2C**; PERMANOVA R^2^=0.35, p=0.016). Communities from the upper monimolimnion (21 m, 24 m) separated from those of the lower monimolimnion (30 m, 45 m, 52.5 m) along the first principal coordinate axis (**Figure S2C**), and the communities at 21 m and 24 m were more similar to each other than to any other depth (**Figure 3B**, **Figure S2D**). To determine whether the biogeochemical gradients in FGL (**Figure 2A**, **Table S1**) were correlated with microbial community composition, we used constrained correspondence analysis (CCA). We found that microbial community composition was strongly correlated with sulfide (CCA ANOVA F=22.71, p=0.001) and methane (CCA ANOVA F=8.07, p=0.003), and marginally correlated with ammonia concentrations (CCA ANOVA F=2.69, p=0.06) (**Figure 3A**). Concentrations of dissolved organic carbon (DOC) were not correlated with bacterial community structure (CCA F=2.48, p=0.09). While depth explained only 35% of the variation in community structure in the PERMANOVA model, the first two PCoA axes (**Figure S2B**), which may represent biogeochemical and physical gradients such as light, accounted for 86% of the variation in community structure.

**Figure 3.**
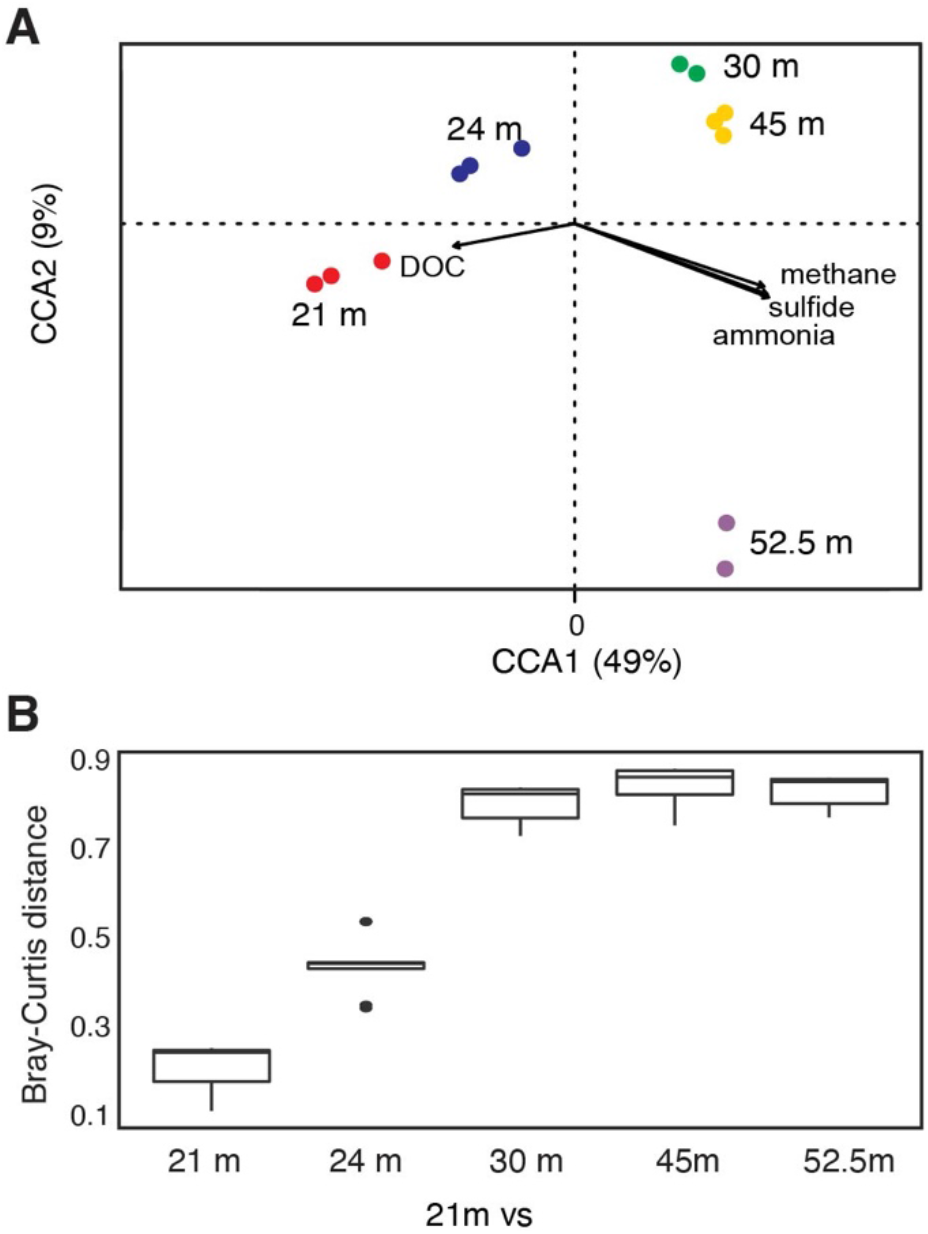
FGL biogeochemistry drives microbial community structure along the water column. (A) CCA analysis correlating bacterial community structure at the sampled depths with geochemical gradients. Only the two primary CCA axes are shown and samples are color-coded by depth. Arrows indicate the direction and magnitude of the relationship (ANOVA: sulfide F=22.71, p=0.001; methane F=8.07, p=0.003; ammonia F=2.69, p=0.06; DOC F=2.48, p=0.09). (B) Bray-Curtis dissimilarity between 21 m and all other sampled depths (n for each depth = 3, except for 30 and 52.5 m where n=2).

### 3.3 Organic electron donors are strong drivers of microbial community assembly in enrichment cultures

To identify and examine microbial interactions and processes involved in the cycling of carbon, sulfur, and other elements below the chemocline, FGL sediments were inoculated into sterile media that contained different combinations of organic electron donors and terminal electron acceptors. The organic electron donors were selected to reflect the diversity of possible organic electron donors in FGL, whereas the terminal electron acceptors represented the most common electron acceptors available in anoxic, sulfidic environments. To identify major sources of variation among enrichment cultures, we used PERMANOVA and PCoA analysis. These analyses revealed the clustering of bacterial communities from enrichment cultures by simple (e.g., acetate, propionate) versus more complex (e.g., chitin, cellulose, lignin) polymers along PCoA Axis 1 (**Figure 4A**, **Table S3**). Subsequent statistical analyses supported this finding; organic electron donors explained 54% of the variation in community structure across samples (**Figure 4A**; PERMANOVA R^2^=0.542, p=0.001). By comparison, terminal electron acceptors only explained 10% of the observed variation (PERMANOVA electron acceptor R^2^=0.104, p=0.001).

**Figure 4.**
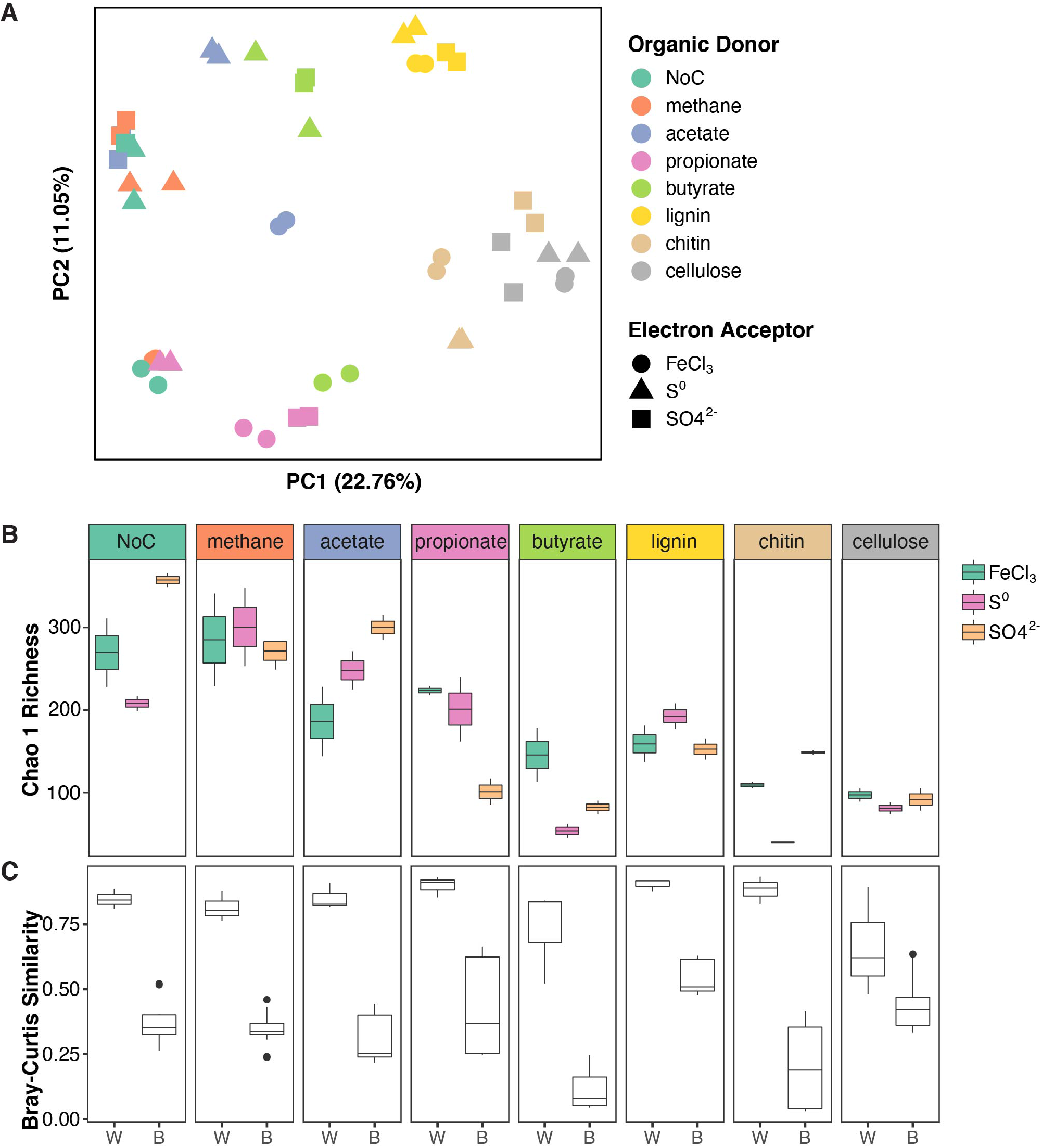
Structure and diversity of enrichment cultures grown on different organic electron donors and terminal electron acceptors. (A) PCoA of enrichment cultures based on Bray-Curtis distances. Samples are color coded based on the organic electron donor (i.e. carbon source) supplied and shapes indicate the terminal electron acceptor. (B) Boxplots of Chao 1 richness in enrichment cultures. The boxplots are plotted for each organic electron donor and are color-coded by electron acceptor. (C) Boxplots showing Bray-Curtis similarity values within enrichment cultures (same organic electron donor and terminal electron acceptor) and between enrichment cultures (same organic electron donor and different terminal electron acceptor).

The evenness of bacterial communities from enrichment cultures varied with both organic electron donor and the terminal electron acceptor supplied (linear model LRT for Shannon diversity: organic electron donor χ_7_^2^ =49.27, p<0.001; terminal electron acceptor χ_2_^2^=16.99, p<0.001; interaction term χ_2_^2^=15.04, p<0.001). Bacterial community richness also varied with electron acceptor, but only in enrichments supplied with specific organic electron donors (linear model LRT for Chao 1 Richness: organic electron donor χ_7_^2^ =33.59, p<0.001; terminal electron acceptor χ_2_^2^=1.93, p=0.16; interaction term χ_2_^2^=4.25, p<0.001). Generally, microbial communities in enrichments that were amended by the polymers cellulose, chitin and lignin were less diverse than the communities that utilized other organic electron donors (**Figure 4B**, see **Table S6** for post-hoc comparisons). Additionally, enrichments amended with SO_4_^2−^ were more diverse compared to the enrichments grown in the presence of other terminal electron acceptors (**Table S6**). Microbial community similarity within replicates, i.e., cultures grown on the same organic electron donor/terminal electron acceptor pair was greater than the similarity between conditions, i.e., cultures grown on the same electron donor with different terminal electron acceptors (**Figure 4C**, ANOVA on Bray-Curtis distances: “within” versus “between” F=238.82, p<0.0001).

The type of terminal electron acceptor [Fe(III), SO_4_^2−^, S^0^] only accounted for 10% of the observed variation in community structure across samples, but the interaction between organic electron donors (i.e., carbon source) and terminal electron acceptors explained an additional 32% of the variation (PERMANOVA interaction term R^2^=0.321, p=0.001). Specifically, the bacterial community composition in enrichment cultures that had been amended with methane, acetate, butyrate, or no carbon source exhibited a stronger dependence on the terminal electron acceptors and larger spread along PCoA Axis 2 (**Figure 4A**) compared to the cultures amended with propionate, lignin, chitin or cellulose.

Organic electron donors selected for specific microbial taxa (**Figure 5**, **Figure S3, Table S7**). Bacterial communities in enrichment cultures amended by methane and propionate, or those that were not amended by any organic electron donors, were similar in composition and were dominated by anaerobic respiring genera *Acetobacterium (*Eubacteriaceae), *Desulfocapsa* (Desulfocapsaceae), *Desulfurivibrio* (Desulfurivibrionaceae), *Desulfomicrobium* (Desulfomicrobiaceae; in NoC and methane), *Desulfobulbus* (Desulfobulbaceae; in propionate only) (**Figure 5**, **Figure S3**). The hydrogen-utilizing and acetate-producing *Acetobacterium* had the highest relative abundances in unamended enrichments, and in those amended with methane or acetate (**Figure 5**, **Figure S3, Table S7**). Given that *Acetobacterium* spp. can ferment carbon substrates such as glucose, fructose and lactose, generating acetate as a waste product (Balch et al., 1977), this organism likely produced the acetate in enrichments that were not amended by any organic electron donors. This likely fueled the growth of sulfur and sulfate reducers. The former organisms include *Geobacter*, which was only enriched on acetate and chitin enrichments supplemented with S^0^, and the latter include *Desulfomicrobium, Desulfocapsa, and Desulfurivibrio*. In fact, the communities amended by acetate were composed exclusively of *Geobacter* and unclassified taxa from Geobacteraceae.

**Figure 5.**
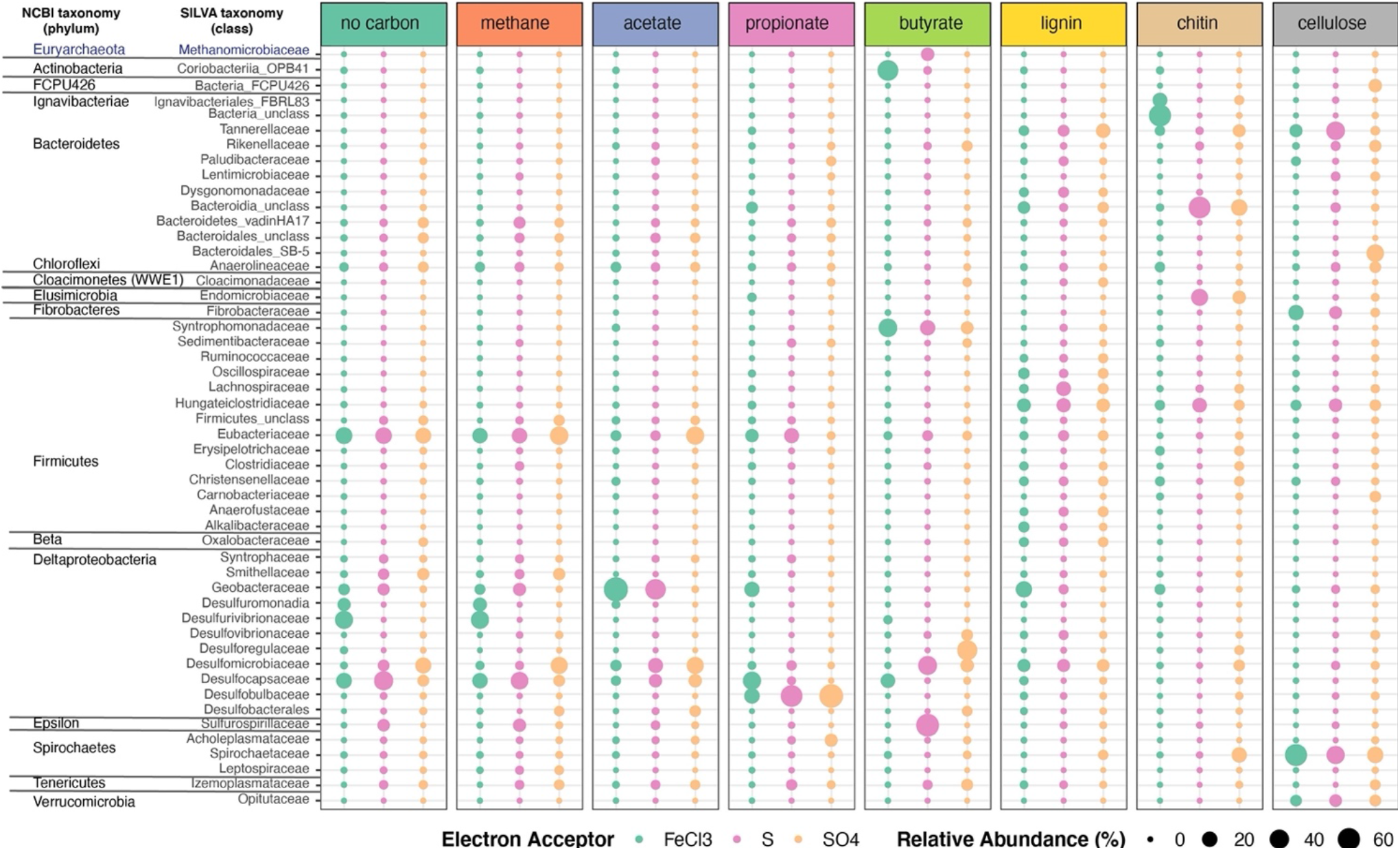
Microbial taxonomic profiles of enrichment cultures from FGL sediments. Plots show the proportion of 16S rRNA gene sequences assigned to each bacterial family across enrichment cultures grown under different organic electron donors and terminal electron acceptors. Only bacterial taxa >0.23% abundances are shown. Profiles are color-coded by the terminal electron acceptor supplied and larger bubbles indicate higher relative abundances.

Bacterial communities that grew on chitin, cellulose or lignin mainly contained fermenting bacteria. These communities did not resemble each other or those grown on other carbon sources (**Figure 5**, **Figure S3**). Enrichment communities amended with chitin and iron comprised the largest abundances of one ASV that was classified as unclassified Bacteria (55%). This ASV was 100% identical to the sequences of uncultured bacteria from aquatic environments (Suzuki et al., 2013) and 97.5% identical to *Cytophaga xylanolytica* (Yarza et al., 2013), a known fermenter of mono-, di-, and polysaccharides except cellulose. Additional taxa present in chitin and Fe(III) enrichments included Ignavibacteriales from the phylum Firmicutes (15.6%), and Geobacteriaceae (4.3%) from the Desulfobacteriota phylum (**Table S7**). Chitin enrichments amended with sulfur or sulfate contained *Endomicrobium* sp. from the phylum Elusimicrobiota (26% and 11.5%) and Sphingobacteriales (1.7%) (**Table S7**). In chitin amendments supplied with sulfur, the dominant ASV was classified as unclassified Bacteroidia. Other abundant taxa in enrichments amended by chitin and Fe(III) were *Endomicrobium* sp. (26%) from the phylum Elusimicrobiota, *Pseudobacteroides* sp. (12.5%; Firmicutes), and Reikenellaceae (1.5%). It is not known whether any of these organisms are also capable of anaerobic respiration using sulfur or sulfate as terminal electron acceptors. The communities amended with chitin and Fe(III) contained low amounts of organisms capable of anaerobic respiration. Enrichments amended by lignin contained the same unclassified ASV as enrichments amended with chitin, as well as several different genera from Firmicutes (*Acetobacterium*, *Anaerofustis*, Christensenellaceae, and Lachnospiraceae), Bacteroidota (Dysgonomonadaceae), and Macellibacteroides (Porphyromonadaceae). Enrichments amended with cellulose harbored abundant fermenting anaerobes such as *Treponema* (Spirochaetaceae), Macellibacteroides (Porphyromonadaceae) and Bacteroidales (**Figure 5**, **Figure S3**). Fibrobacteraceae and Fibrobacter (15.7%) were found enriched in amendments with cellulose and Fe(III) but were not well represented in other enrichments (**Table S7**). Although not abundant, some organisms such as Marinilabiliaceae were only found in enrichments amended with cellulose and lignin.

The composition of enrichment cultures also varied markedly with the terminal electron acceptor supplied. Butyrate-grown communities amended with SO_4_^2−^ harbored abundant sulfate reducing *Desulforegula* (Desulforegulaceae), whereas those amended by S^0^ were dominated by *Sulfurospirillum* sp., a Campylobacterales (**Figure 5**, **Table S7**). These organisms can typically reduce sulfur, thiosulfate and polysulfide, nitrate or manganate, but are unable to reduce sulfate (Goris and Diekert, 2016) (**Figure 5**, **Table S7**). The diversity of sulfate reducers in the enrichments varied with both terminal electron acceptors and organic electron donors. Sulfate reducing *Desulfomicrobium*, *Desulfobulbus* and *Desulforegula* spp., preferred acetate, propionate, and butyrate, respectively (**Figure 5**). Propionate is commonly used to isolate sulfate reducing microbes (Samain et al., 1984; Sorokin et al., 2012); indeed, this substrate enriched *Desulfobulbus* and Desulfocapsaceae. Cultures amended with Fe(III) harbored *Geobacter* (and Geobacteraceae*), Desulfurivibrio* spp, and *Desulfocapsaceae* spp., which are capable of dissimilatory iron reduction (Kuever et al., 2005).

Members from the phylum Euryarchaeota and Halobacterota (both domain Archaea) were retrieved from all enrichments albeit at small relative abundances (<~1% for most enrichments) (**Figure S3**, **Table S7**). These organisms, particularly Halobacterota, had the highest relative abundances in the cultures grown on butyrate and S^0^. In addition, we identified putative methanogens and methanotrophs based on the classification provided by Evans et al. (2019) and Smith and Wrighton (2019) at low abundances in all enrichments. Putative methanotrophs were most abundant (<0.5%) in enrichments that contained methane or no organic electron donor. Notably, similarly low abundances of methanotrophs were recovered from the monimolimnion of FGL, and included members from Micrarchaeales and Woesearchaeales (**Table S4**).

### 3.4 Network analysis elucidates potential microbial interactions involved in the cycling of carbon, sulfur, and other compounds

To identify potentially significant syntrophic (positive) or competitive (negative) interactions among microbial groups, we utilized network analyses of bacterial taxa from different depths in the lake and from enrichment cultures of the sediment (see Methods). The microbial network from the upper monimolimnion contained 179 nodes (taxa) with 19.8 edges per node on average, whereas the network from lower monimolimnion was less dense and had 148 nodes with 4.9 edges per node (Table S8, **Figure 6**). Nodes classified as Chlorobi, *Sulfuricurvum* sp. and sulfate reducing bacteria (such as *Desulfocapsa*, *Desulfatiglans*, *Desulfobacca,* and *Syntrophus*) represented “hub” taxa that connected different microorganisms in the upper monimolimnion. Some of those nodes connected to >20 other taxa (the average number of edges = 14.7, Table S8). In the deeper waters, sulfur-reducing bacteria (e.g., Desulfocapsa), sulfur-oxidizing (Syntrophus (Syntrophaceae)) and fermenting organisms appeared to facilitate connections among microbes: these ASVs shared positive interactions with other community members in the lower monimolimnion (**Figure 6**). For example, several members from the Campylobacteraceae (*Sulfurovum* and *Sulfurimonas*) and Desulfoarculaceae, Desulfosarcinaceae, *Desulfocapsa* (Desulfobacterota, prev. Deltaproteobacteria) had positive interactions with members from the Bacteroidales and Chloroflexi groups.

**Figure 6.**
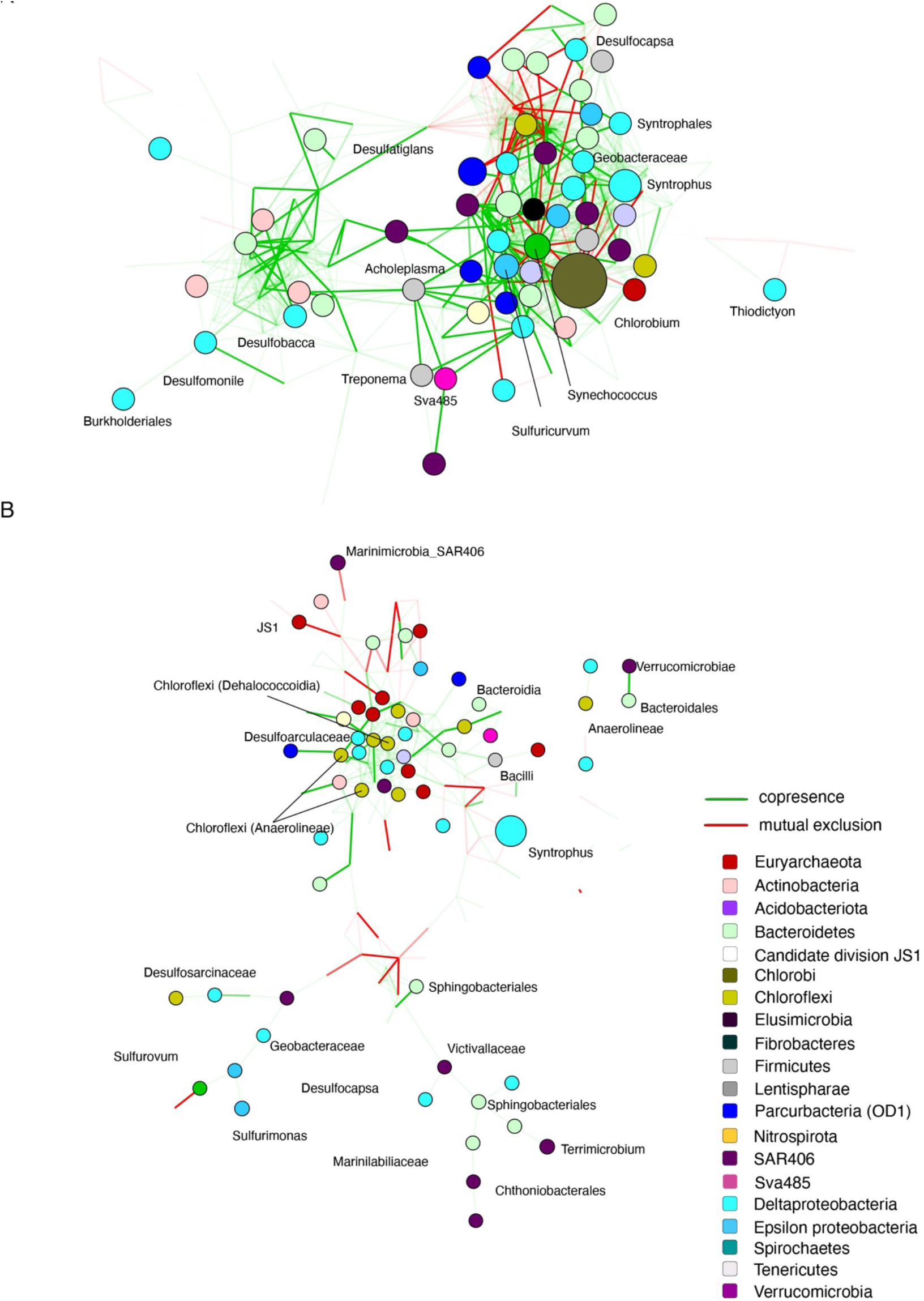
Significant co-occurrence and co-exclusion relationships among bacterial taxa in the FGL water column. Networks of microbial communities from the (A) upper monimolimnion (21 m and 24 m) and (B) lower monimolimnion (30 m, 45 m, and 52.5 m). Networks were built from 16S rRNA gene profiles using CoNet in Cytoscape. Nodes represent taxa (genus level), and are colored by phylum. Each edge represents a significant co-occurrence/mutual exclusion relationship (p<0.05; green=co-occurrence, red=exclusion).

Several mutually exclusive, competitive interactions were also observed among taxa from the lake. In both the upper and lower monimolimnion, there was a mutually exclusive relationship between members of Deltaproteobacteria (e.g., Desulfobacterales), and Marine Group A bacteria (SAR406) (**Figure 6**). Marine Group A bacteria lack cultured representatives, but these organisms have been implicated in the cycling of sulfur in marine environments. Their genomes harbor genes for polysulfide reductase, an enzyme necessary for dissimilatory polysulfide reduction to hydrogen sulfide, and for dissimilatory sulfur oxidation (Wright et al., 2014).

Microbial interactions observed in the anaerobic water column and in enrichments supplemented with cellulose and lignin suggested the involvement of organisms such as *Macellibacteroides*, Paludibacteraceae and members from Bacteroidales and Bacteroidia groups. in the degradation of plant material below the chemocline (Table S4, Table S7, **Figure S4**). Microorganisms that were enriched on S^0^ or Fe(III) were present in networks from the lower monimolimnion, indicating downward transport of microbes and oxidants from the chemocline to this region. For example, microbes enriched on iron oxyhydroxide and organic polymers (e.g., chitin, lignin and cellulose) were present in the monimolimnion, and included Sphingobacteriales (up to 1.9% in cultures amended with chitin and SO_4_^2−^), Ignavibacteriales (~1%), and Anaerolineaceae (up to 5%; **Table S4**). Co-occurrence analysis indicated a positive relationship among these microorganisms and *Desulfobacterium* sp. (**Figure 6** and **Figure S4**). Microbial networks from the lower monimolimnion included members from the class Dehalococcoidia (Chloroflexi). These organisms contain *dsr* genes, are likely capable of sulfite reduction (Wasmund et al., 2014; Wasmund et al., 2016), and can obtain energy from the oxidation of substituted homocyclic and heterocyclic aromatic compounds (Löffler et al., 2013). Several sulfate reducers, also capable of dissimilatory iron reduction, such as *Desulfobacter* sp., *Desulfobacterium* sp., and *Desulfobulbus* sp. (Lovley, 2006) were found both in the lower monimolimnion and enrichment cultures supplied with cellulose, lignin or propionate amended by any of the electron acceptors.

### 3.5 Bacterial communities in FGL contain a high number of bacterial specialists and few bacterial generalists

Our analyses supported niche partitioning of bacterial communities around specific organic compounds. Certain microbial groups in enrichment cultures were rare and found almost exclusively in the presence of particular electron donors or acceptors (e.g., Endomicrobium/Endomicrobiaceae in chitin-amended cultures; **Figure 5**). Others were ubiquitous and present across many conditions (e.g., Desulfomicrobiaceae, Desulfocapsaceae, Eubacteriaceae; **Figure 5**). This suggested that bacterial generalists and specialists were present in both the water column and enrichment cultures. To quantify the bacterial specialists and generalists, we calculated Levin’s niche breadth, which revealed that the water column and enrichment samples harbored more generalists than specialists **(Figure S5).** The generalists included Anaerolineales, Bacteroidia, Desulfobacteria, Gammaproteobacteria, Syntrophia (Desulfobacterota) and two Archaea (Nanoarchaea, and Thermoplasmata), among others. Bacterial taxa that were present across enrichment cultures and the water column included Anaerolineales, Bacteroidales, Campylobacterales, Izemoplasmatales (Bacilli), and Syntrophales (Desulfobacterota). According to Levin’s niche breadth, bacterial specialists in the water column belonged to Burkholderiales (upper monimolimnion), Dehalococcoidia (at 45 and 52.5 m), Micrococcales (at 45 m) and Sphingobacteriales (coincidentally also present in chitin and cellulose cultures) (**Figure S5**). The taxa Desulfarculales, Desulfobulbales (methane and no carbon cultures), Desulfuromonadia (no carbon), Methanobacteriales (butyrate), and Oscillospirales (lignin) were classified as bacterial specialists in enrichment cultures supplemented by specific organic electron donors. These experimental conditions and observations pointed to the degradation of the plant material, particularly of lignin and cellulose, throughout the anoxic water column of FGL.

## 4 DISCUSSION

### 4.1 Comparisons of microbial community profiles in FGL to those found in other meromictic lakes

The microbial communities in the monimolimnion of FGL were vertically stratified and likely shaped by geochemical and physical gradients. Prior studies of FGL reported anoxygenic anaerobic phototrophic purple-sulfur bacteria (PSB) as the most abundant clones at 20.5 m (25%) and GSB as less abundant (15%) (Meyer et al., 2011). PSB were not abundant in the upper monimolimnion (at 21 m and 24 m) at the time of our sampling. Given that our filtered samples from 21 m were purple, the 16S rRNA gene primers used in the amplification of the DNA from our samples likely selected against this group. In any case, the recovery of 16S rRNA sequences of anoxygenic phototrophs (green sulfur bacteria, purple sulfur bacteria and cyanobacteria) from sediments (data not shown) provided evidence for the vertical transport of organic matter from the chemocline to the lower monimolimnion and sediments. Anoxygenic phototrophs and sulfate reducers at or immediately below the chemocline (upper monimolimnion) are abundant in many meromictic and monomictic lakes (Klepac-Ceraj et al., 2012; Baricz et al., 2014; Andrei et al., 2015; Llorens-Mares et al., 2015; Baatar et al., 2016; Meyerhof et al., 2016; Kadnikov et al., 2019).

In keeping with previous studies of sulfidic lakes, we found that microbial community richness and evenness increased with depth: communities within and closer to the chemocline were less diverse compared to those from the deeper waters. Reduced diversity is also reported at the oxic/anoxic interface of Ursu Lake (Andrei et al., 2015) and Mahoney Lake (Klepac-Ceraj et al., 2012), and in the upper layers of meromictic Lake Oigon (Baatar et al., 2016). Similarly, the highest species richness was observed in the monimolimnion of six meromictic lakes in the Palau region (Meyerhof et al., 2016). The low bacterial diversity at the chemoclines of FGL and other meromictic lakes that contain sulfide within the photic zone stems from the overabundance and blooms of a few taxa of anoxygenic phototrophic bacteria (Klepac-Ceraj et al., 2012; Baricz et al., 2014; Andrei et al., 2015; Llorens-Mares et al., 2015; Baatar et al., 2016; Meyerhof et al., 2016; Kadnikov et al., 2019).

Microbial community composition in the monimolimnion of FGL was strongly correlated with concentrations of sulfide and methane, and modestly correlated with ammonia concentrations, suggesting a tight coupling between the microbial community composition and biogeochemical cycling in this meromictic lake. The compositions of bacterial communities of lakes Shunet, Shira, and Oigon were also strongly correlated with sulfide concentrations (Baatar et al., 2016). Although we did not measure light availability throughout the water column, this physical factor likely drives the microbial community structure in the upper monimolimnion. Light availability and intensity are known to influence the microbial composition and function in microbial mats and the photic zones of sulfidic lakes (Hartgers et al., 2000; Franks and Stolz, 2009; Diao et al., 2017). In FGL, spectral analyses show that light does not penetrate past the base of the chemocline (Brunskill and Ludlam, 1969). The known adaptations of GSB to low light environments may explain why they are the dominant bacterial group at 21 m and 24 m (Overmann et al., 1992; Borrego et al., 1999; Marschall et al., 2010).

Our study expanded on 16S rRNA gene surveys of environmental samples by establishing targeted enrichment cultures and conducting network analyses to assess the contributions of specific microbial taxa, organic electron donors and inorganic electron acceptors to biogeochemical cycling in the lake. Specifically, comparisons of the microbial communities at different depths of the lake to enrichments amended with pairs of terminal electron acceptors and organic electron donors typically found in this lake allowed us to assess the functional potential of microorganisms and their interactions in the monimolimnion. The anoxic and sulfidic deeper waters of FGL harbored more diverse communities compared to the chemocline. These communities contained many candidate phyla with no known cultured representatives, collectively termed “microbial dark matter” (Rinke et al., 2013). These candidate phyla included Caldatribacteriota (OP9/JS1), Patescibacteria (OD1), Dependentiae (TM6), Elusimicrobiota and others (**Table S4**); their 16S rRNA sequences and/or metagenomes have been recovered from several meromictic lakes (Borrel et al., 2010; Comeau et al., 2012; Klepac-Ceraj et al., 2012; Gies et al., 2014). The relative abundance of candidate phyla with no known cultured representatives increased with depth in FGL and these taxa co-occurred with sulfate reducers. Our enrichment culture experiments and the available metagenomes of Caldatribacteriota (OP9/JS1) are consistent with these organisms fermenting carbohydrates and hydrocarbons (Dodsworth et al., 2013; Nobu et al., 2016; Liu et al., 2019) and producing hydrogen and acetate that can be used by methanogens (Hug et al., 2013; Wrighton et al., 2014). Other abundant taxa in the lower monimolimnion included Chthoniobacterales (Verrucomicrobiae), Syntrophus (Desulfobacterota) and Sphingobacteriales. Chthoniobacterales were also present in the cultures amended by cellulose, which was consistent with their noted preference for carbohydrates, and the capability for the degradation of cellulose, chitin, and xylan (Cabello-Yeves et al., 2017). Members of Verrucomicrobiae have been recently found to contain some of the earliest evolved *dsrAB* genes involved in sulfite reduction (Anantharaman et al., 2018). These genes are also present in dissimilatory sulfate or sulfite reducers like Desulfobacterota (Wagner et al., 1998) and are likely involved in the cycling of sulfur in addition to degrading carbohydrate polymers.

### 4.2 The availability of organic electron donors drives microbial community structure in FGL

Our findings highlighted electron donors as strong drivers of the microbial community assembly in the enrichment cultures; this predictor explained 58% of the variation in community structure across samples compared to 10% explained by terminal electron acceptors (Figure 4). No study thus far has evaluated the combined effects of individual terminal electron acceptor and organic electron donor on the assembly of microbial communities. Kwon et al. (2016) supplied acetate, lactate or glucose to anaerobic microcosms and reported that the microbial communities were structured by these organic electron donors. However, these communities were grown in enrichments amended with both sulfate and ferrihydrite electron acceptors. Chen et al. (2017) investigated the oxidation of acetate, propionate, and butyrate in enrichments amended with both nitrate and sulfate and found that nitrate was reduced first with a preference for propionate and acetate, sulfate reduction followed and was coupled to the oxidation of butyrate. These observations bolster the support for interactions between organic electron donors and terminal electron acceptors as strong determinants of community structure in stratified environments, and suggest microbial preference for specific electron donor/terminal electron acceptor pairs. Our experiments expand the understanding of how numerous donor-acceptor pairs influence microbial assembly and demonstrate the roles of the diversity and availability of organic electron in structuring microbial communities in the monimolimnion.

Various lines of evidence in our study and previous studies show that organic matter is transported from the chemocline to the lower monimolimnion and sediments. This particulate organic matter likely includes plant and zooplankton material from the oxic zone and aggregates of photosynthetic microbes and minerals from both the oxic and anoxic zones (Thompson and Ferris, 1990; Thompson et al., 1997; Hubeny et al., 2011; Meyer et al., 2011; Fulton et al., 2018). Indeed, microorganisms that were enriched on lignin, cellulose and chitin such as Bacteroidales Ignavibacterales, Elusimicrobia, and Anaerolineaceae were also detected in the lower monimolimnion. We predict that many of the microbes capable of degrading polymers are associated with the sinking particles in the FGL and that particle-associated populations are likely enriched in the hydrolytic/fermentative specialists that we observed in treatments amended with complex carbohydrates such as lignin, cellulose and chitin. Conversely, microorganisms enriched on DOM such as acetate, butyrate or propionate are more likely to be enriched in the free-living (planktonic) fraction. Differences in the microbial community composition and metabolic potential between the free-living and particle-associated communities have been observed in several environments, and microbes associated with the particles had a higher richness and abundance of functional genes involved in carbon degradation, nitrogen fixation, and stress response than the free-living microbes (Lemarchand et al., 2006; Fuchsman et al., 2011; Garcia et al., 2013; Liu et al., 2020). Thus, we hypothesize that particle-associated bacteria actively contribute to the biogeochemical cycling in FGL.

### 4.3 Possible contributions of microbial interactions and bacterial generalists and specialists to the cycling of carbon, sulfur and iron in FGL

Our findings demonstrate a first-order resemblance of FGL microbial communities to the microbial communities in other sulfidic meromictic lakes and provide novel evidence for the extensive cross-feeding interactions among microbes. We propose that these interactions contribute to the high levels of microbial diversity observed in the monimolimnion and are key to the cycling of carbon in anoxic aquatic environments.

Co-occurrence is more frequently observed among groups of organisms that share functional traits or have cross-feeding partnerships (Freilich et al., 2011; Zelezniak et al., 2015). In networks from both the water column and enrichment data, dissimilatory sulfate reducing bacteria such as Desulfobulbaceae or Desulfobacteraceae co-occurred with strictly anaerobic fermenting and hydrocarbon-consuming Anaerolineaceae (Chloroflexi) (Liang et al., 2015). Similar cross-feeding interactions between fermenting bacteria and sulfate or sulfur reducers have been observed in mixed cultures amended with chitin and cellulose (Bharati et al., 1982; Pel et al., 1989). Other microorganisms associated with Synthrophaceae in the upper monimolimnion belonged to Verrucomicrobia and the sulfate reducing Desulfoarculaceae. In enrichment cultures amended with organic polymers (cellulose, chitin, or lignin), the network analysis also showed positive interactions of Syntrophaceae with the methane producing members of the phylum Euryarchaeota (**Figure S4**). Overall, these results suggest that the syntrophic interactions with Synthrophaceae are critical to the oxidation of organic carbon in the lower monimolimnion. Importantly, some of the interactions observed in the FGL, such as those that depend on photosynthetic activity and the presence of oxygen, are likely driven by biogeochemical and physical gradients and not strictly biological interactions such as cross-feeding.

The generalist bacterial taxa found at all depths in the FGL water column (Desulfatiglandales, Synthrophales, Sphingobacteriales) have the capacity for sulfate reduction (Jørgensen, 1982), an essential metabolism of many aquatic environments and sediments (Jørgensen and Kasten, 2006; Jørgensen et al., 2019). Conversely, bacterial specialists were only recovered from certain depths or enrichment conditions. An example of these are members from Endomicrobium genus. These microbes were not detected in the lake water column, but were enriched in cultures amended with chitin and either sulfate or sulfur, but not iron. Based on our enrichments, these microbes can digest chitin. The only known cultured representative of this group, *Endomicrobium proavitum*, was isolated from a termite gut and can fix nitrogen (Zheng et al., 2016). However, this microbe is not a known chitin degrader. Microorganisms enriched on cellulose, such as *Treponema* sp. and Macellibacteroides were not very abundant in other enrichment cultures and were found in trace amounts in the lower monimolimnion. The microbial processes involved in the anaerobic decomposition of cellulosic material are limited by electron acceptor availability and require diverse syntrophic bacterial interactions, which might explain why these metabolisms are not more widespread among bacteria (Bao et al., 2019). In general, the concentrations and the types of available and labile electron donors, including organic polymers, may be limited in deeper waters. The degradation of these donors may require specific bacterial groups that produce intermediates that are then consumed by other microbial taxa, giving rise to a diverse microbial community in the monimolimnion (Seth and Taga, 2014).

Syntrophic metabolisms and cross-feeding interactions are integral to the cycling of carbon and sulfur in many anoxic environments (Schink, 1997; Overmann and Schubert, 2002; Sieber et al., 2012) and several syntrophic groups were present in FGL. In the lower monimolimnion, the high relative abundances of Syntrophaceae (>30% at 30 m; >40% at 45 m) (**Table S4**) suggested syntrophic interactions, possibly with methanogens or some other hydrogen-utilizing microorganisms. In the enrichments, Syntrophaceae were not very abundant (typically <2%). The highest enrichments, 2.8 %, were observed in cultures with no added carbon or with methane. Although we did not recover putative methanogens from the upper or the lower monimolimnion by 16S rRNA gene primers, in the metagenomes from 45.5 and 52 m, methanogens had a relative abundance of 3.5% at 45.5 m % and 3.1% at 52 m. Additionally, archaeal taxa potentially capable of synthesizing or metabolizing methane were recovered from both the water column (~3.5%, e.g. Woesearchaeales) and enrichments (<1%; e.g. Methanomicrobiales). Lastly, we also observed positive interactions between Syntrophaceae and Verrucomicrobia or Bacteroides. It is possible that Verrucomicrobia and Bacteroides ferment hydrocarbons and carbohydrate polymers and provide organic acids as organic electron donors to Syntrophaceae to couple with the reduction of sulfate.

### 4.4 Outlook

The composition of microbial communities in the anoxic and sulfidic zones of meromictic lakes is tightly coupled to the physical and biogeochemical gradients. Understanding the functions of individual microbial members and their interactions can better inform predictions of the effects of climate and anthropogenic changes on carbon sequestration in these lakes. Our study provides a framework for investigating the contributions of microbial communities to nutrient cycling in meromictic lakes. Future analyses should be conducted at a finer spatial scale and should examine microbial community diversity in different size fractions to determine the influences of planktonic and particle-associated microbial communities in nutrient cycling. Furthermore, in order to test hypotheses regarding the influence of specific organic compounds and physical processes that distribute the organic matter on microbial community structure, organic carbon substrates such as acetate, cellulose, and amino acids should be characterized throughout the monimolimnion. Analyses of metagenomes and metatranscriptomes of bacterial communities and the development of more comprehensive reference databases of environmental microorganisms will be critical for furthering our understanding of the microbial ecology of these lakes. Lastly, detailed characterization of microbial community succession in enrichments amended with different electron acceptors and donors represent a fruitful avenue for identifying the specific microbes and microbial interactions that are crucial for nutrient cycling in meromictic lakes.

## AUTHOR CONTRIBUTIONS

TB and VKC designed the study; AdST, SP, and VKC collected samples; CR processed lake samples; AdST prepared and SP processed all enrichments; CR, AFS, SP, and VKC analyzed the sequence data; CR, VKC, and TB wrote, edited and finalized the manuscript. All authors approved the final version of the manuscript.

## ACKNOWLEDGMENTS

We thank Shikma Zaarur, David Wang and Sharon Newman for helping out with sample collection, and to Libusha Kelly and Kevin Bonham for helpful comments on the manuscript. The project was funded by Simons Foundation (TB), Wellesley College Fiske grant (VKC) and NIH UG3 OD023313 (VKC).

## DATA AND CODE AVAILABILITY

Raw amplicon sequence reads for both the water column samples and enrichment cultures were deposited in NCBI’s Sequence Read Archive (SRA), under BioProject PRJNA613903 and accession numbers SAMN14480503-SAMN14480563. Unless otherwise noted, plots were generated in R v.3.6.2. Annotated R scripts for conducting the statistics and figures showed in this manuscript are available on Github: https://github.com/rojascon/FGL_2021_FrontiersinMicrobiology.

## SUPPLEMENTARY MATERIAL

The Supplementary Material (figures and tables) for this manuscript can be downloaded online at bioRxiv, alongside this submission.

