## Supplementary Figures for "Organic electron donors and terminal electron acceptors structure anaerobic microbial communities and interactions in a permanently stratified sulfidic lake"

**Figure S1. Reproducible abundances and variation of bacterial taxa along the FGL watercolumn. A)** Rarefaction curves for FGL 16S rRNA gene sequences grouped at 97% sequence identity. **B)** FGL bacterial communities at each sampled depth (21 m, 24 m, 30 m, 45 m and 52.5 m). Only genera with average relative abundances >1% are shown, and all others were clumped into Other.

**A)**

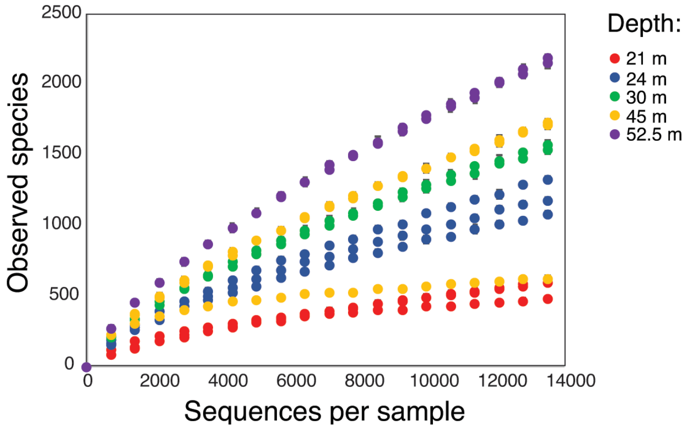


**B)**

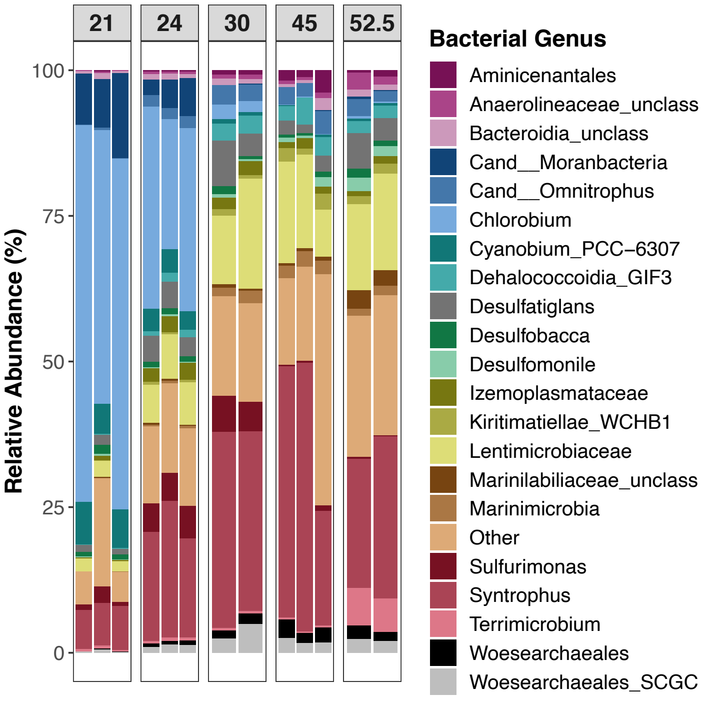

**Figure S2. Vertical stratification of bacterial communities along the FGL water column**. Stacked bar plots showing the relative frequency of 16S rRNA gene sequences assigned to each bacterial A) phylum and B) order across samples. Samples are grouped by depth, and each color represents a bacterial order. C) PCoA plots constructed from Bray-Curtis dissimilarity matrices. Each point represents a sample and is color-coded by depth. Closeness of points indicates high community similarity. The % of variance accounted for by each principal-coordinate axis is shown in the axis labels. D) Average Bray-Curtis dissimilarity values among samples from different depths. Long boxplots indicate high variability in pairwise dissimilarity values.

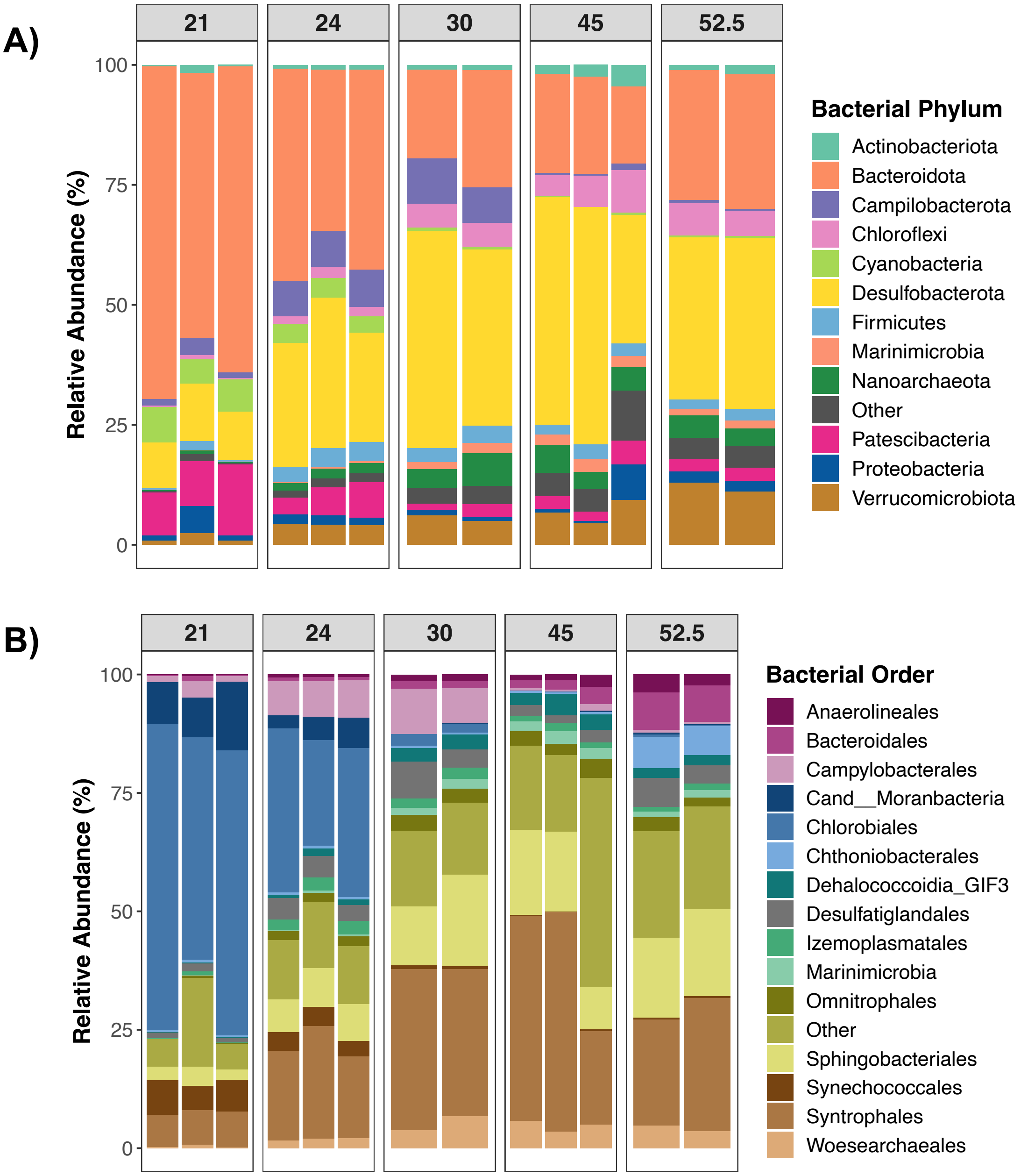

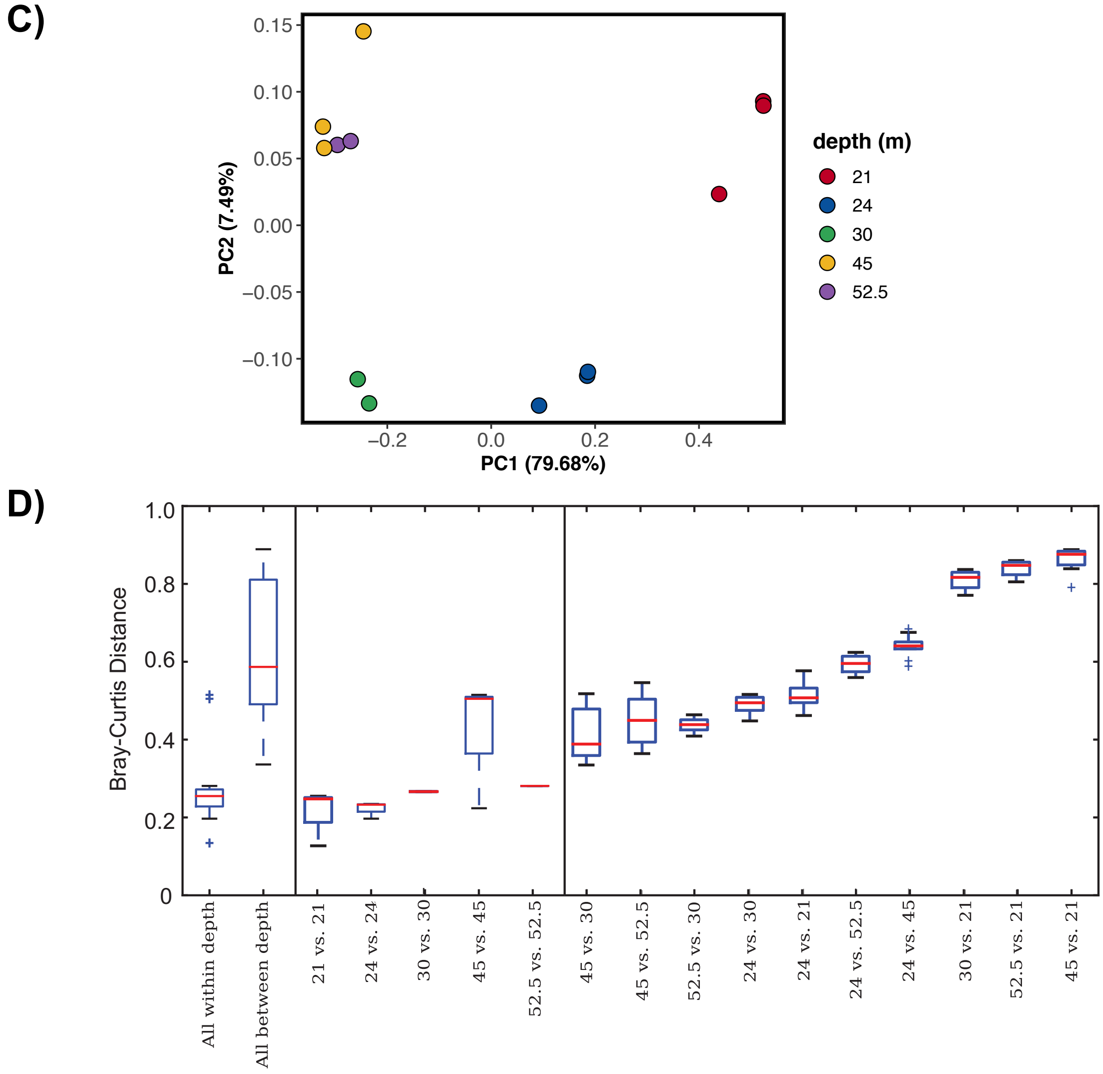
C)
0.15
0.10
0.05
0.00
−0.05
−0.10
PC2 (7.49%)
−0.2
0.0
0.2
0.4
PC1 (79.68%)
depth (m)
21
24
30
45
52.5
D)
Bray-Curtis Distance
1.0
0.8
0.6
0.4
0.2
0
All within depth
All between depth
21 vs. 21
24 vs. 24
30 vs. 30
45 vs. 45
52.5 vs. 52.5
45 vs. 30
45 vs. 52.5
52.5 vs. 30
24 vs. 30
24 vs. 21
24 vs. 52.5
24 vs. 45
30 vs. 21
52.5 vs. 21
45 vs. 21

**Figure S3. Predominant bacterial taxa of enrichment cultures amended with different organic electron donors.** Stacked bar plots showing the relative frequency of 16S rRNA gene sequences assigned to each bacterial A) phylum, B) order , or C) genus across samples. Samples are grouped by the organic electron donor (i.e. carbon source) supplemented, ordered by electron acceptor ($FeCl_3$, $Na_2SO_4$, $S^0$) and each color represents a bacterial phylum/order. Only phylum/orders with average relative abundances >1% are shown, and all others were clumped into Other.

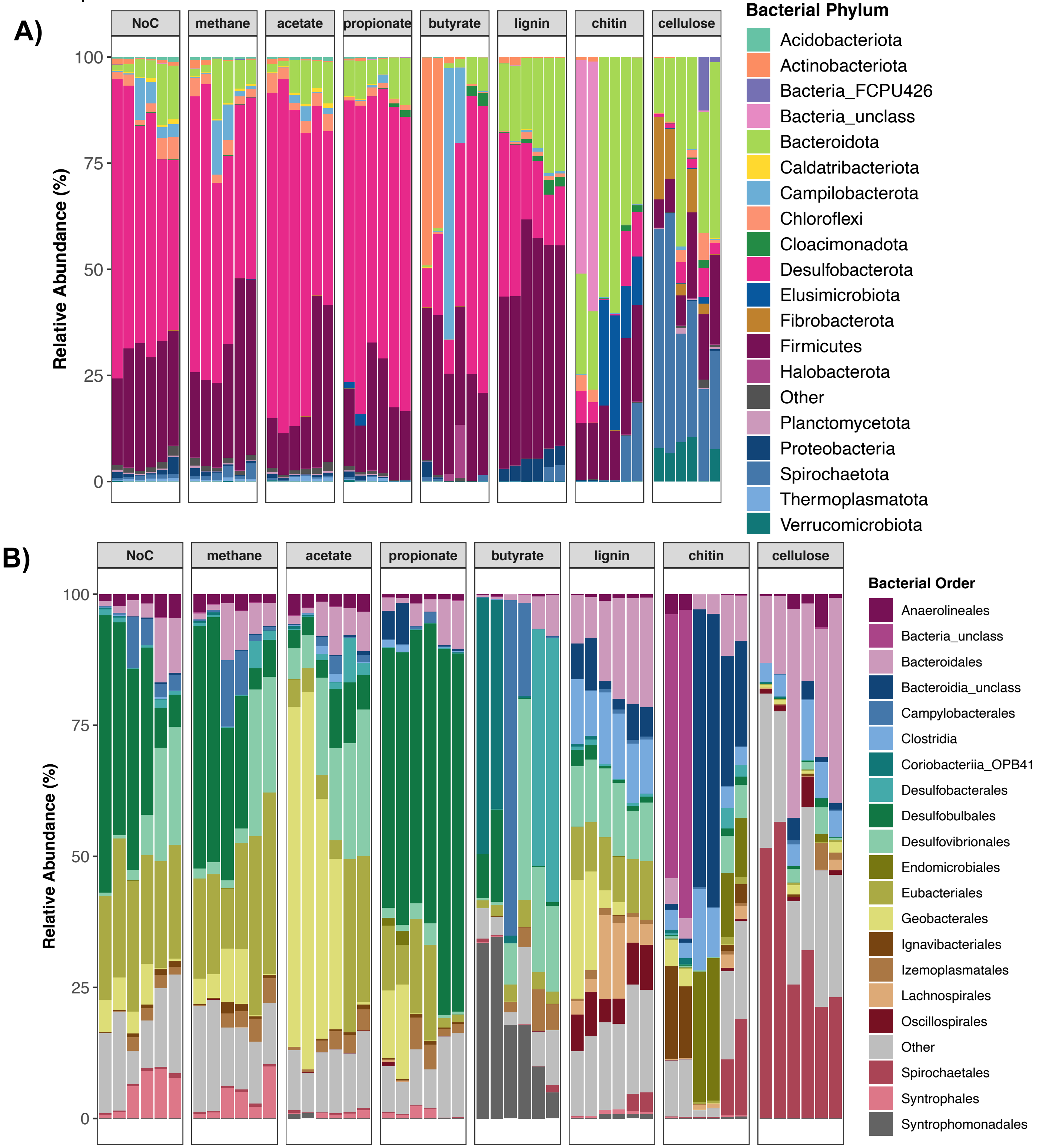

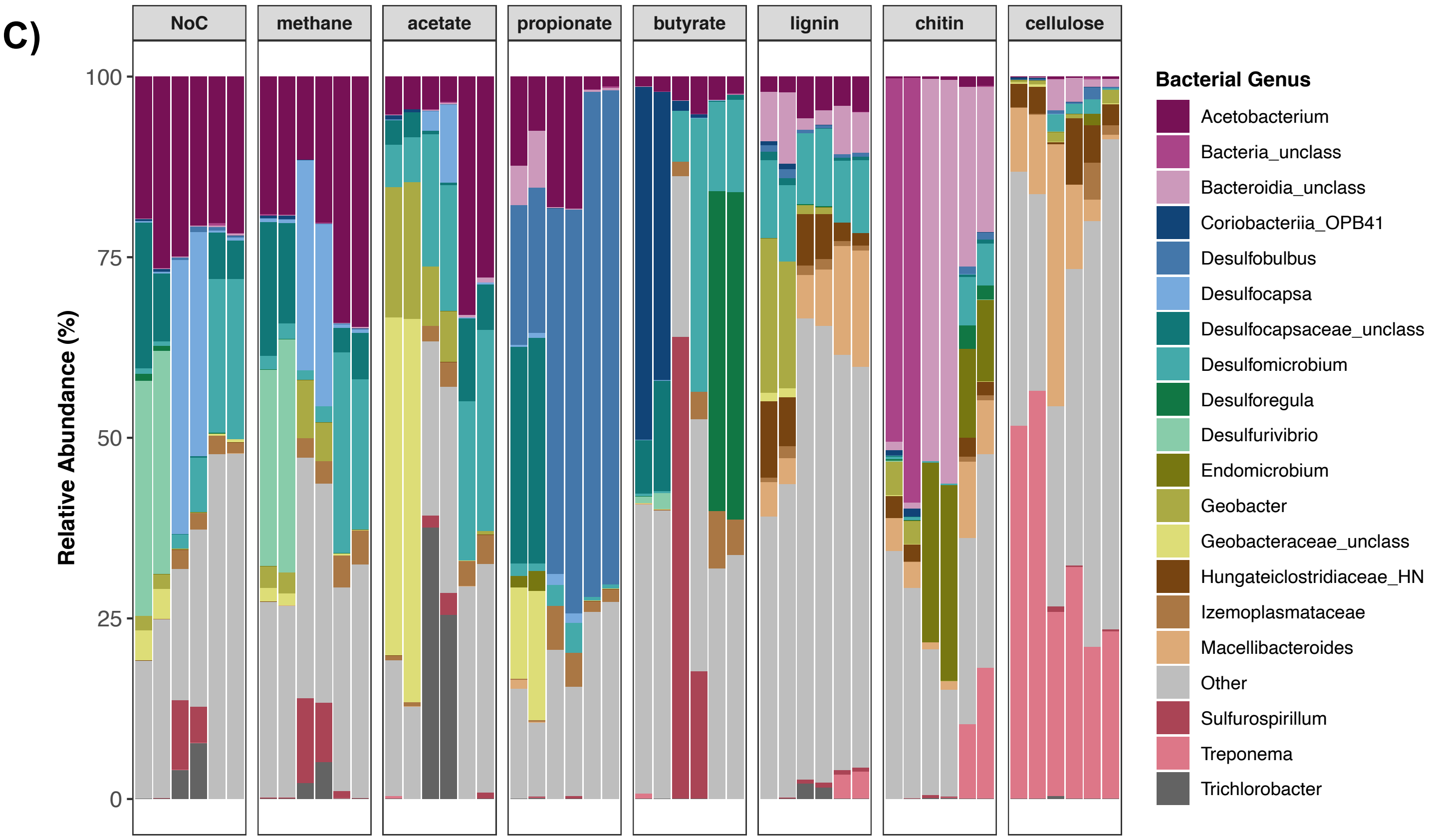

C)
NoC
methane
acetate
propionate
butyrate
lignin
chitin
cellulose
Relative Abundance (%)
100
75
50
25
0
Bacterial Genus
Acetobacterium
Bacteria_unclass
Bacteroidia_unclass
Coriobacteriia_OPB41
Desulfobulbus
Desulfocapsa
Desulfocapsaceae_unclass
Desulfomicrobium
Desulforegula
Desulfurivibrio
Endomicrobium
Geobacter
Geobacteraceae_unclass
Hungateiclostridiaceae_HN
Izemoplasmataceae
Macellibacteroides
Other
Sulfurospirillum
Treponema
Trichlorobacter

**Figure S4. Significant co-occurrence and co-exclusion relationships among bacterial taxa in enrichment cultures from FGL sediments.** Microbial community networks of enrichment cultures supplemented with different terminal electron acceptors (Fe, S°, $SO_4$). Networks were built form 16S rRNA gene profiles using CoNet in Cytoscape. Nodes represent bacterial taxa (genus level), and are colored by phylum. Each edge represents a significant co-occurrence/mutual exclusion relationship ($p<0.05$; green=co-occurrence, red=exclusion).

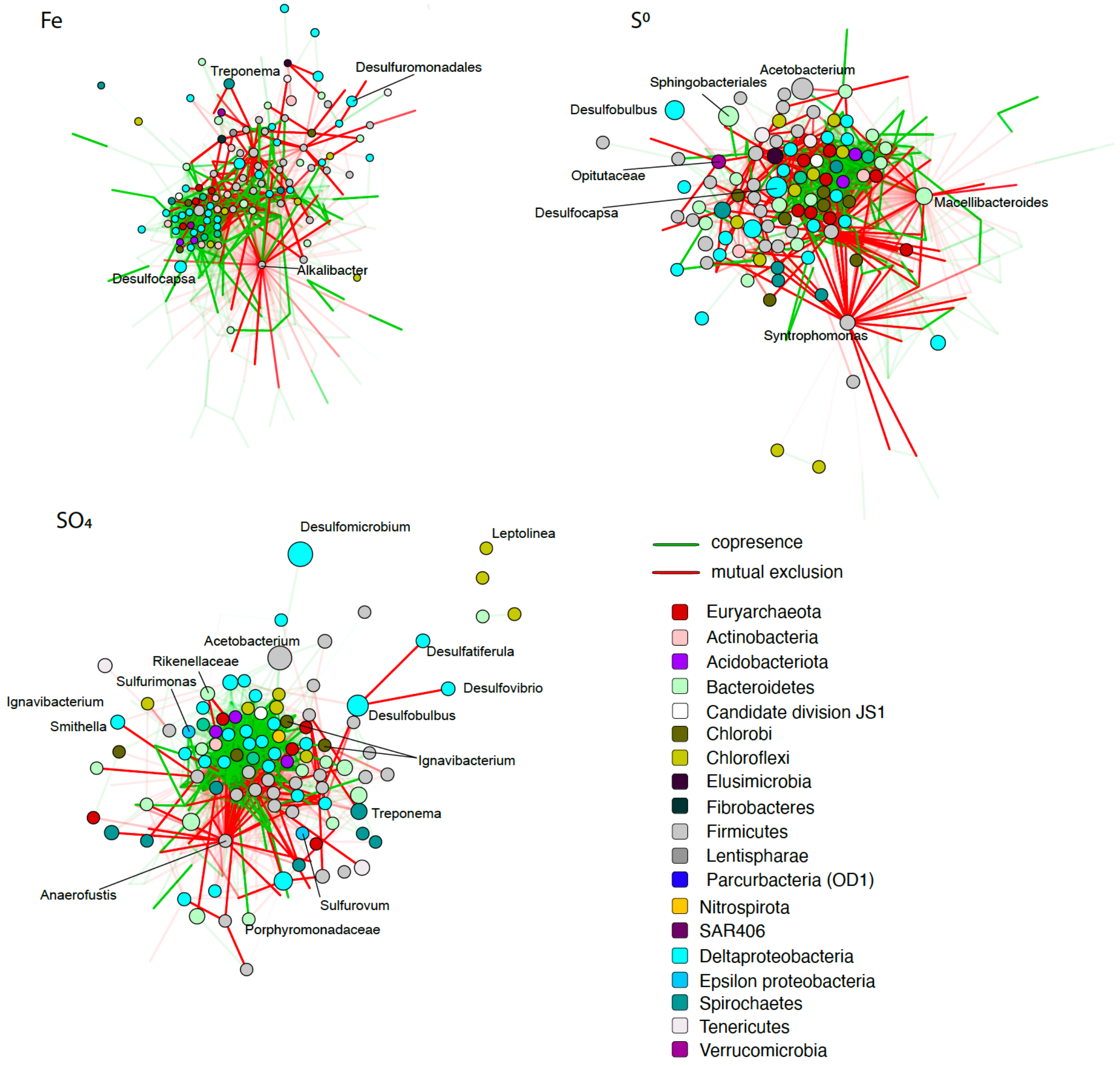

**Figure S5. Bacterial ASVs classified as generalists or specialists in FGL bacterial communities.** Plots of Levin's niche breadth vs. average relative abundance (log) for bacterial ASVs from (A) water column samples and (B) enrichment cultures. ASVs of interest are color-coded by their bacterial order; and dashed lines indicate ±1 SD of the mean niche breadth. Note how the color-coding is not identical between the two plots.

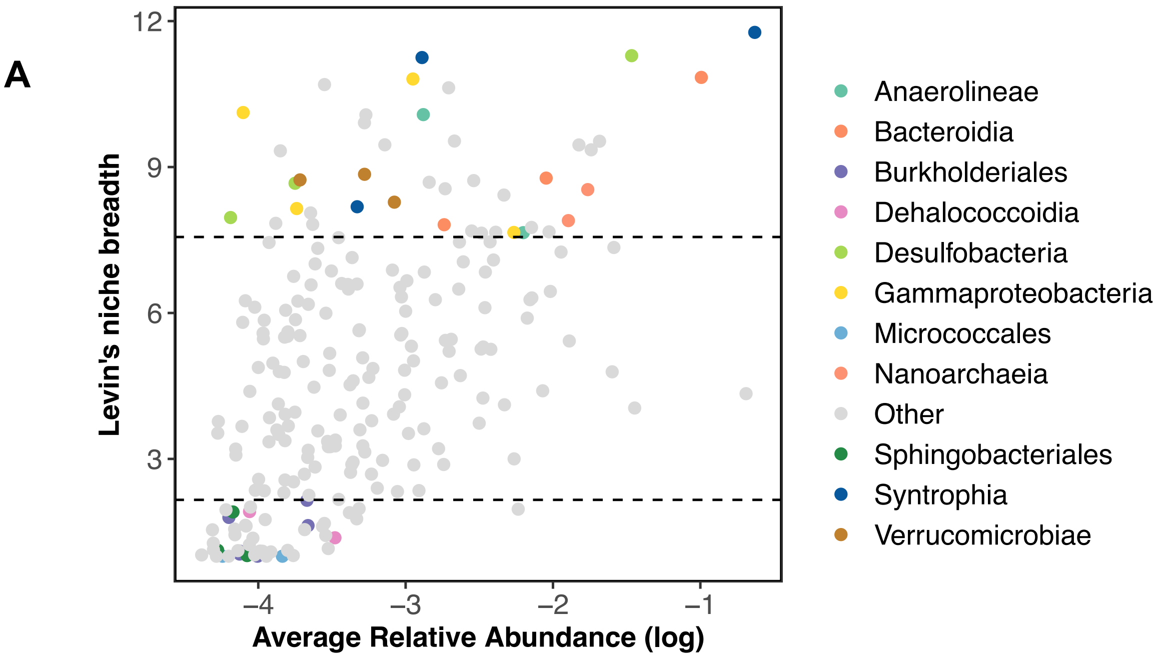


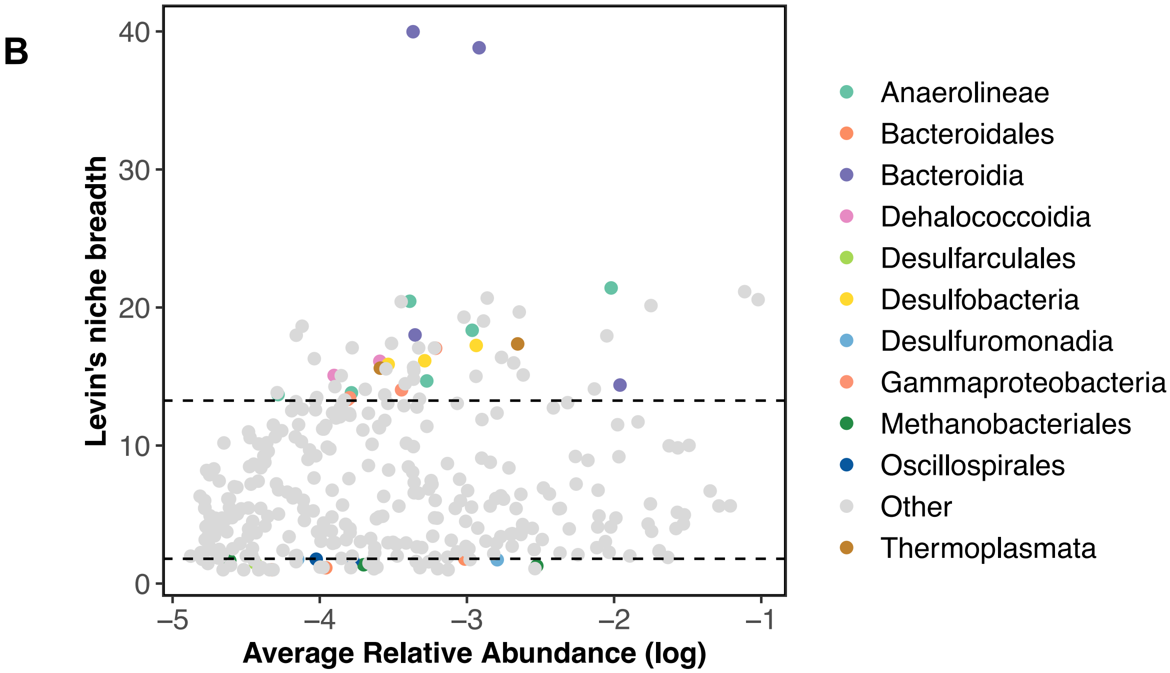
